# Spatiotemporal dynamics of molecular expression pattern and intercellular interactions in glial scar responding to spinal cord injury

**DOI:** 10.1101/2021.12.20.473346

**Authors:** Leilei Gong, Yun Gu, Xiaoxiao Han, Chengcheng Luan, Xinghui Wang, Yufeng Sun, Mengya Fang, Shuhai Yang, Lai Xu, Hualin Sun, Bin Yu, Xiaosong Gu, Songlin Zhou

## Abstract

Adult regeneration in spinal cord is poor in mammalian but remarkable in the neonatal mammals and some vertebrates, including fish and salamanders. Increasing evidences basis of this interspecies and ontogeny highlighted the pivotal roles of neuron extrinsic factors-the glial scar, which exert confusing inhibiting or promoting regeneration function, but the spatiotemporal ordering of cellular and molecular events that drive repair processes in scar formation remains poorly understood. Here, we firstly constructed tissue-wide gene expression measurements of mouse spinal cords over the course of scar formation using the spatial transcriptomics (ST) technology in Spinal cord injury (SCI) repair. We analyzed the transcriptomes of nearly 15449 spots from 32 samples and distinguished normal and damage response regions. Compared to histological changes, spatial mapping of differentiation transitions in spinal cord injury site delineated the possible trajectory between subpopulations of fibroblast, glia and immune cell more comprehensively and defined the extent of scar boundary and core more accurately. Locally, we identified gene expression gradients from leading edge to the core of scar areas that allow for re-understanding of the scar microenvironment and found some regulators in special cell types, such as Thbs1 and Col1a2 in macrophage, CD36 and Postn in fibroblast, Plxnb2 and Nxpe3 in microglia, Clu in astrocyte and CD74 in oligodendrocyte. Last, we profiled the bidirectional ligand-receptor interactions at the neighbor cluster boundary, contributing to maintain scar architecture during gliosis and fibrosis, and found GPR37L1_PSAP and GPR37_PSAP were top 2 enriched gene-pairs between microglia and fibroblast or microglia and astrocyte. Together, the establishment of these profiles firstly uncovered scar spatial heterogeneity and lineage trajectory, provide an unbiased view of scar and served as a valuable resource for CNS injury treatment.

**Highlights:**

- Spatial illustration of gene expression pattern after T10 right lateral hemisection.
- Spatial atlas of scar formation by 21 cell types around damaged area.
- The origin, trajectory reconstruction and functional diversity of cell types in different stages of scar formation.
- Novel insights for glial scar boundary and potential benefits for recovery intervention after SCI.

## Introduction

Spinal cord injury (SCI) mostly caused by accidents is a lifelong, traumatic injury with a tremendous social and economic impact (*1, 2*). More worryingly, the clinical therapeutic methods are too limited to promote spinal cord regeneration and the outcomes are complicated closely related on the level of the injury (*3, 4*). The pathophysiological processes of SCI involve a primary injury and the formation of a glial scar (*5*). Although it is widely accepted that the glial scar can inhibit axon regeneration and protect spared neural tissues by injury, the exact role of the glial scar in SCI remains poorly understood (*6*). During the approximately 2 weeks, the glial scar becomes mature which associated with many cellular and extracellular components and their complex interaction. The cellular components of glial scar mainly consist of macrophage, fibroblast, microglia, astrocyte, oligodendrocyte, NG2^+^ oligodendrocyte precursor cells (NG2-OPCs) and ependymal cells. The extracellular matrix (ECM) includes chondroitin sulfate proteoglycans (CSPGs), myelin-associated glycoprotein (MAG), fibronectin, oligodendrocyte myelin glycoprotein (OMGP), matrix glycoprotein tenascin C and hyaluronan fragments (*7–9*). To date, the related researches of cellular mechanisms and molecular regulation of these components are still in progress (*10*). Therefore, it is essential for getting a comprehensive understanding of above events to develop more effective treatments inducing disability for SCI.

Rapid advancements in single-cell RNA sequencing (scRNA-seq) technologies have permitted the use of single cell transcriptional profiling to investigate cellular heterogeneity within the CNS, advancing our knowledge of CNS disease. For example, astrocytes, a major cell type found throughout the CNS, have a wide array of functions in the modulation of synaptic transmission, regulation of blood flow, blood-brain barrier formation and the metabolic support of other brain resident cells, yet the character of their cellular heterogeneity and how it manages these diverse roles remains unclear. Using a fluorescence-activated cell sorting-based strategy, five distinct astrocyte subpopulations were identified across three brain regions that show extensive molecular diversity. These populations differentially support synaptogenesis between neurons. Specially, the emergence of specific subpopulations during tumor progression corresponded with the occurrence of seizures and tumor invasion (*11*). Additionally, five distinct astrocyte subtypes were identified in adult mouse cortex and hippocampus (*12*). Recently, a population of disease-associated astrocytes was identified in an Alzheimer’s disease mouse model using scRNA-seq. These disease-associated astrocytes appeared at early disease stages and increased in abundance with disease progression (*13*). Single-cell transcriptomics also revealed that the human ganglionic eminence depends more heavily on intermediate progenitor cells as workhorses than does the developing neocortex, with its greater reliance on radial glial cells (*14*). Moreover, by using scRNA-Seq, intra-glial scar gene expression heterogeneity and interactions have been documented at the level of individual cells (*15*).

A common feature to transcriptome-wide scRNA-seq studies is that they do not address the spatial patterns of gene expression. Although the computational inference can partially circumvent the lack of spatial information for scRNA-Seq data (*16, 17*), we still need alternative methods that can provide positional information on the different cell types involved in the process, which is necessary for a comprehensive understanding of glial scar dynamics (*8, 18*). Although current in situ sequencing techniques, such as expansion-assisted iterative fluorescence in situ hybridization (EASI-FISH), have been exploited (*19–23*), these approaches have limitations. It requires prior knowledge about the genes of interest, and uses padlock probes and rolling circle amplification in tissue sections to target known genes. The biggest flaw is that the approach can only detect a small numbers of marker genes (*19, 24*). The spatial dimensions of entire transcriptomes remain unexplored in glial scar, along with the spinal cord micro-environment.

Spatial transcriptomics (ST) is an untargeted technology that produces quantitative transcriptome-wide RNA sequencing data and determines the quantitative spatial distribution of polyadenylated transcripts in a tissue section using barcoded oligo-dT DNA capture probes arrays corresponding to histological imaging (*25*). The throughput of the approach is substantially greater than that of other spatially resolved methods. It produces spatial maps based on RNA-seq data and has broad applicability (*26–28*). For example, studies on the brain (*25, 29, 30*) the spinal cord of neurodegeneration(*31*) and the development of human heart (*32, 33*) have offered spatial cell-specific findings that could not have been accomplished using traditional tissue homogenates. It contributes to confirm regional markers and cell type identifications based on scRNA-seq and ST.

Here we applied ST to spatially profile gene expression and cell types in thoracic (T10) spinal cord tissue sections from WT mice to present a spatiotemporal atlas that systematically describes the spatial archetypes and cellular heterogeneity of the scar formation at three stages in the early acute stage (3 dpi), subacute stage (7 dpi) and intermediate stage (14 dpi, 28 dpi) after spinal cord injury (Figure 1A). We obtain the unique molecular signatures of multiple cell types as well as their subpopulations and pseudotime trajectory throughout the acute and chronic injury stage. By assessing expression of bidirectional ligand-receptor pairs and spatial distribution on different subpopulations, we identified scar architecture and re-understanded of the scar micro-environment. As the first ST analysis of spinal cord scar, our transcriptomic dataset aims to present a comprehensive decoding of the glial scar and explore the potential therapeutic strategies in clinical.

**Figure. 1.**
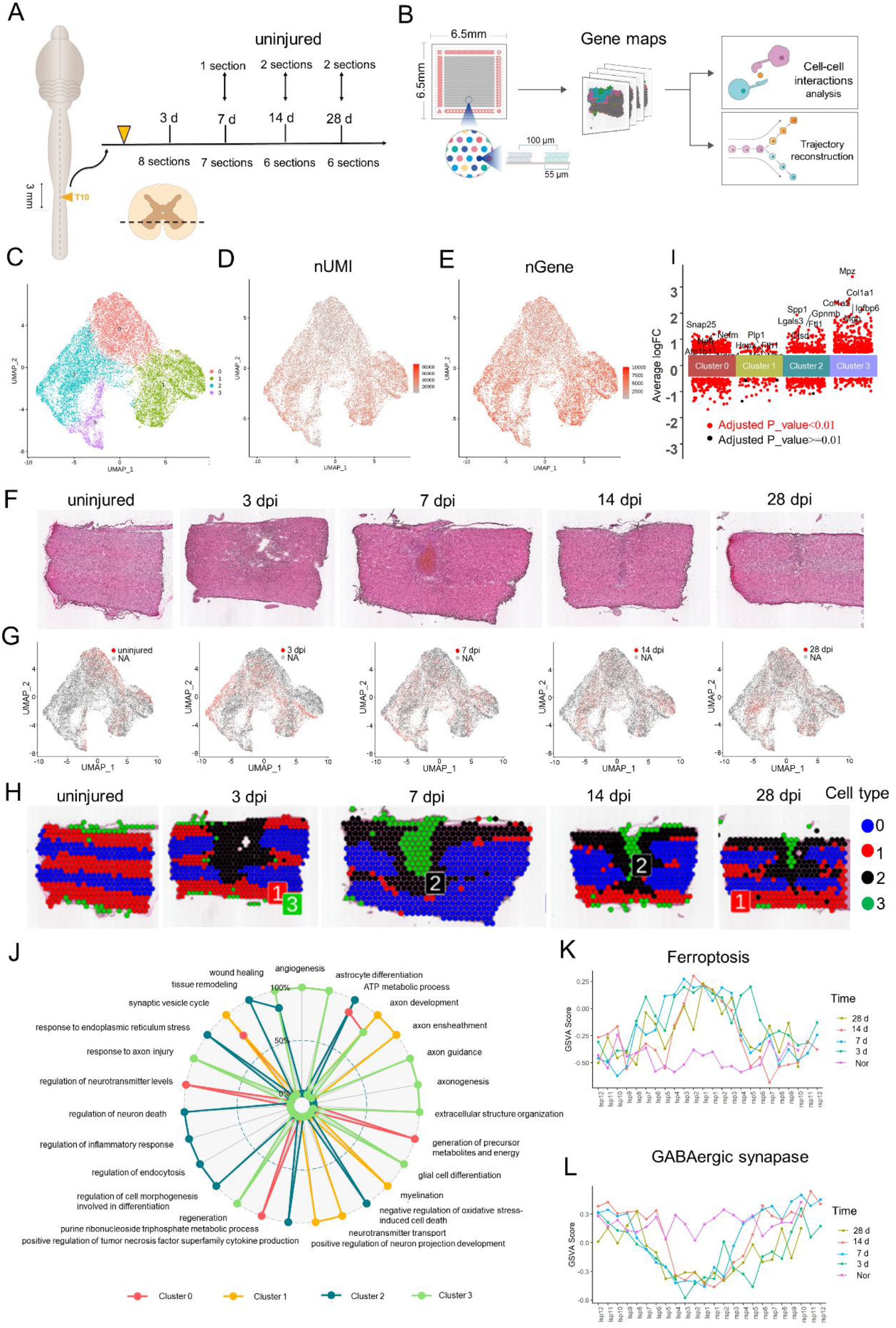
Generation of a spatiotemporal transcriptional atlas of mouse glia scar. (A) Overview of the whole study design for glia scar formation and location of sections used in this study. (B) Experimental workflow and analysis for spatial RNA-seq of glia scar in adult mouse at four stages of scar maturation after T10 right lateral hemisection. Spatial microarrays have 4992 spatially barcoded spots of 55 µm diameter and 100 µm center-to-center distance. The Spatial Transcriptomic (ST) procedure yields matrices with read counts for every gene in every spot, which are then decomposed by a set of factors (“cell types”). (C) Uniform Manifold Approximation and Projection (UMAP) plot of spots from all the sections was visualized using Seurat (v4.0) package and profiled the cell clusters. (D and E) UMAP profiled the number of the expressed UMI (nUMI) and genes (nGene), respectively. (F) H&E staining images show the changes in pathological morphology of glia scar maturation. (G) UMAP profiled the spots at different time after injury, respectively. (H) The UMAP spots were mapped to their spatial location, respectively. (I) The top 5 highest differentially expressed genes (DEGs) were listed in each cluster listed. (J) The Gene Ontology (GO) analysis data were enriched for each cluster. (K) GSVA score of ferroptosis signaling pathway were counted at three stages in Layer 3. (K) GSVA score of GABAergic synapse pathway were respectively counted at three stages in Layer 3.

## Materials and Methods

### Mice and surgeries

Sixty female mice (eight-week-old, C57BL/6J) were randomly divided into five groups (n=12). The mice were anaesthetized by isoflurane. The procedure for T10 right lateral hemisection was similar to that described previously (*34*). Briefly, a midline incision was made over the thoracic vertebrae. A T10 laminectomy was performed. To produce a right lateral hemisection injury, tips of optimum knife (BVI Beaver, Oakville, Canada) were carefully inserted into posterior median sulcus of the spinal cord then to gently scrape it across the bone on the ventral side to not spare any tissue ventrally or laterally. The muscle layers were sutured, and then the skin was secured with wound clips. The mice were placed on soft bedding on a warming blanket held at 37 °C until fully awake and given ibuprofen for pain relief. All the animal experimental were conducted in accordance within situational animal care guidelines and approved ethically by the Administration Committee of Experimental Animals, Jiangsu Province, China, 20150304-004.

### Spatial transcriptomic tissue imaging, library preparation, and sequencing

Libraries were prepared using the Spatial Transcriptomic Library Preparation Glass Slides following manufacturer’s instructions. In brief, frozen samples were embedded in OCT (TissueTek, Sakura) and stored at −80 °C until cryosection. For cryo-sectioning, samples were equilibrated to −22 °C and were cut into 10 μm thickness using Leica CM3050 S cryostat. Sections were placed on chilled Visium Tissue Optimization Slides (3000394, 10X Genomics) and Visium Spatial Gene Expression Slides (2000233, 10× Genomics), and adhered by warming the back of the slide. Tissue sections were then fixed in chilled methanol and stained with Hematoxylin and Eosin Y according to the Visium Spatial Gene Expression User Guide (CG000239 Rev D, 10× Genomics). For sequencing, tissue was permeabilized for 24 min, which was selected as the optimal time based on tissue optimization time course experiments. Libraries were prepared according to the Visium Spatial Gene Expression User Guide. cDNA synthesis was carried out overnight using SuperScript III Reverse Transcriptase in the presence of Actinomycin. Libraries were loaded at 300 pM and sequenced on a NovaSeq 6000 System (Illumina) using a NovaSeq S4 Reagent Kit (Illumina). Sequencing was performed using the following read protocol: read 1, 28 cycles; i7 index, 10 cycles; i5 index, 10 cycles; read 2, 90 cycles.

### Spatial transcriptomics analysis

Reads were mapped against the Mouse Genome (mm10) and Gencode (vM23) transcripts were quantified using spaceranger-1.2.0 (10× Genomics). The number of genes with at least one UMI count in any tissue covered spot. The combined data were normalized with Seurat 3.2.2. We computed principal components (PCs) to perform dimensionality reduction processing on the combined data, and selected the top 30 principal components for downstream data analysis. We performed graph-based clustering across all samples and displayed by UMAP. Marker genes of the five clusters were analyzed by Seurat 3.2.2. The marker gene function enrichment the Gene ontology (GO) and the Kyoto Encyclopedia of Genes and Genomes (KEGG) pathway analysis in each cluster were performed by cluster Profiler package in R.

### Gene Set Variation Analysis (GSVA)

The pathway activities of selected spots were quantified by applying GSVA package (*35*). In order to show the expression of different genes in space, we integrated the physical coordinates of 4992 points on the spatial transcriptome chip with the coordinate information of the corresponding spot points, and then used the R language ggplot2 package to project them into a two-dimensional space. The UMI value in each spot represented the expression abundance expression (Ei) of each gene. Here, i indicated the expression vector of each spot.

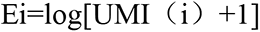

The degree of activation of the signal pathway is reflected in the score value.

### Pseudotime trajectory

Pseudotime trajectories for selected subclusters were constructed using the R package Monocle v2.14. 0(*36*). The raw counts for cells were extracted and normalized by the estimate size factors and estimate dispersions functions with default parameters. Genes detected in more than 50 spots and with an average expression >0.1 were retained. Differential expression analysis was carried out using the differential GeneTest function with a model against the pseudotime. The variable genes with the lowest adjusted q value < 0.001 were clustered and plotted in the heatmap. Cell functions and the trajectory were constructed by the reduce dimension function with default parameters.

### Co-expression network analysis

The first co-expression networks were constructed using weighted correlation network analysis (WGCNA) (v1.66) package in R (*37*). After fltering genes, gene expression values were imported into WGCNA to construct co-expression modules using the automatic network construction function. TOM Type is unsigned and minModule Size is 8. Genes were clustered into 14 correlated modules. To find out biologically significant modules, module eigengenes were used to calculate the correlation coefficient with samples or sample traits. Intramodular connectivity and module correlation degree (MM) of each gene were calculated by R package of WGCNA, and genes with high connectivity tended to be hub genes which might have important functions. Correlation analysis was performed using module eigengene with data for specific traits or phenotypes. Pearson correlations between each gene and trait data under the module are also calculated for the most relevant module (positive correlation and negative correlation) corresponding to each phenotype data, and gene significance value (GS) was obtained. Another an unbiased coexpression networks were constructed using the method that described previously (38). The first 2000 highly variable variables in all spots were normalized, and the correlations were calculated by pearson correlation coefficient. The coexpression modules were obtained by k-mean clustering.

### Ligand-Receptor interaction analysis

To infer potential cellular crosstalk between different cell types, we adapted the method used in CellPhoneDB for single-cell transcriptome data (*39*). Enriched receptor–ligand interactions were predicted between two cell types based on expression of a receptor by one cell type and a ligand by another cell type. Only receptors and ligands expressed more than a specified threshold of the cells in the specific cluster are considered significant (default is 0.1). The pairwise comparisons between all neighbor cell types in the scar were then performed. First, we determined the mean of the average receptor expression level in a cluster and the average ligand expression level in the interacting neighbor cluster by randomly permuted the cluster labels of all cells. The p-value for the likelihood of cell-type specificity of a given receptor–ligand complex was obtained by calculating the proportion of the means which are as or higher than the actual mean.

### Immunostaining

Spinal cord sections were first fixed with 4% paraformaldehyde and then incubated overnight at 4°C in the following primary antibodies: Iba1 (Wako,Rabbit 1:100), GFAP (abcam, Chicken 1:100), P4HB (abcam, Mouse 1:100), F4/60 (abcam, Rabbit 1:200), CD68 (NOVUS Biological, Mouse 1:100), Laminin (abcam, Mouse 1:300), Fn1 (LSBio, Sheep 1:100), Collagen IV (SouthernBiotech, Goat 1:100), Thbs1 (Invitrogen, Mouse 1:100), Col1a2 (GeneTex, Rabbit 1:100) and Nxpe3 (abbexa, Rabbit 1:100) diluted in PBS/0.3% Triton/5% normal donkey serum. After washing the section with PBS, secondary Alexa Fluor-conjugated antibodies (Invitrogen) diluted in PBS/0.3% Triton were incubated for 2 h at room temperature. Slides were mounted with mounting medium containing DAPI (Beyotime institute of Biotechnology, Shanghai, China) and kept at 4°C until further microscopic analysis. The sections were visualized under a TCS SP2 confocal microscope (Leica-Microsystems GmbH, Mannheim, Germany) or ZEISS M2 fluorescence microscope (Carl Zeiss Jena GmbH, Jena, Germany) and images from sections were processed in ImageJ.

### Spinal cord dehydration and transparency

Dehydration, immunostaining and clearing of spinal cord were done based on previous studies (*40*). After washing away excess secondary antibodies, tissue was gradually dehydrated in resistant glassware with incremental concentrations of tetrahydrofuran (TFH, 50%, overnight; 80%, 1 h and 100%, 2 h, v/v % in distilled water, Sigma-Aldrich). Incubations were performed on a shaker at room temperature. Then remaining lipids were extracted with dichloromethane for 1 hr and dibenzyl ether (DBE) was used for refractive index matching. Samples were kept in a full vial of DBE and protected from light during the whole process to reduce photo bleaching of the fluorescence. Imagings were processed with lightsheet microscope 18 (LS18) by Nuohai Life Science (Shanghai) Co., Ltd.

## Result

### Overview of investigate the spatial gene expression in T10 spinal cord sections

To ensure the uniformity of the wound area, we adapt the lateral hemisection model, instead of contusion (*41*). Previous study revealed that the region of glial scar approximately distributed 0.5 mm away from the site of injury after SCI (*42*). To decipher the cell type and region of glial scar as much as possible, we harvested the spinal cord tissue 1.5 mm away from the site of T10 right lateral hemisection (figure 1A). The Visium Spatial Gene Expression Slides were used to capture and profile the mRNAs in the spots of spinal cord sectons at 3d, 7d, 14d and 28d after T10 right lateral hemisection, which intended to cover the dynamic process of glial scar formation after injury (*43*). Because each section contained up to 4,992 spots in its capture area and the size of the Visium Spatial Gene Expression Slides is 6.5 mm×6.5 mm (figure 1B), approximately four spinal cord tissues can be placed on the same chip to capture more spots as shown in figure S1A. Supplementary Table S1 contains a summary of the data evaluation. Overall, 15449 spots within the 32 sections were analyzed (Table S2, figure S1A and B). Data showed the median sequencing depth of single spot at approximately 219,442 Unique Molecular Identifiers (UMIs) and 4,875 genes in our study (Table S2, figure S1C). The H&E staining images served as a reference for subsequent unsupervised analysis. We focused on the extent of scar foci and boundary that conduct the scar microenvironment, identification cell type, the reconstruction of cell trajectories, cell-cell interactions within the scar formation (figure 1B).

### Defining the spatiotemporal boundary of scar in the injury site

Integrated clustering analysis of ST from 32 samples using Seurat (v4.0) revealed 4 distinct clusters when visualized on a UMAP plot (figure 1C). These 4 clusters covered all cell types in the SCI site and adjacent uninjured spinal cord. The numbers of UMIs and genes in cluster 1, cluster 2 and cluster 3 were larger than that in clusters 0 (figures 1D and 1E). To better understand the spatial distribution of these cell types, we compared the H&E staining images (figure 1F) with their ST data (figure 1G) and mapped the clusters back to their spatial location (figure 1H). These identified spatially patterns could match the changes in pathological morphology. Specifically, cluster 2 and cluster 3 characteristically distribute in the scar regions showed in H&E staining images, consistent with the larger UMIs numbers. Based on these findings, we inferred that cluster 2 and cluster 3 represented multiple cellular components that are known to comprise the glial scar. On the contrary, cluster 0 and cluster 1 represented cell types in the uninjured spinal cord (figure 1E). Together, we firstly identify the approximate scope of glial scar at three stages.

Meanwhile, we listed the top 5 highest differentially expressed genes (DEGs) in each cluster during this process from ST (figure 1I, figure S1D). The highest DEGs provided a unique molecular signature for each cell type. For example, cluster 0 cells highly expressed typical neuron marker Snap25 and mainly represent the neurons in the gray matter area of the uninjured spinal cord. Cluster 1 cells highly expressed myelin associated protein (such as Plp1, Mbp and Mal) and mainly represent the oligodendrocytes in the white matter area of the uninjured spinal cord.

The highest DEGs in cluster 2 were secreted phosphoprotein-1 (Spp1), glycoprotein nonmetastatic melanoma protein B (Gpnmb), galactose-specific lectin 3 (Lgals3) and cathepsin D (Ctsd). Spp1, also known as OPN (Osteopontin), is a broadly expressed pleiotropic protein (*44*). It has multifunction in the pathophysiology of several inflammatory, degenerative, autoimmune and oncologic diseases (*45*). Spp1 was firstly discovered as a component of bone matrix, but it is also expressed by subsets of neurons including alpha-retinal ganglion cells (αRGC), alpha motor neuron, and several classes of glial cells including astrocyte, Schwann cells, Müller cell and disease-associated microglia (*46*); OPN is an alpha motor neuron marker in the mouse spinal cord (*47*). Recent studies have shown that the levels of Spp1 are affected by neural injury. Spp1 exert both pro- and anti-inflammatory roles and contributes to tissue damage (the Yin) not only by recruiting harmful inflammatory cells to the site of lesion, but also increasing their survival (the Yang) (*45, 48*). For example, Spp1 is capable of stimulating mTOR activity (*49*) and promotes RGC regeneration when introduced in combination with either insulin-like growth factor 1 (IGF-1) or brain-derived neurotrophic factor (BDNF) (*46*). Additionally, Spp1 was reported to stimulate the outgrowth of injured motor axons but not sensory neurons (*50*). Our data showed the spatiotemporal distribution of Spp1 (figure S1E). Spp1 indeed appear on some motor neurons and significantly increased in injury site, consistent with previous studies (*45, 47, 48*). Lgals3 plays a role in numerous cellular functions including cell growth, cell adhesion, apoptosis, pre-mRNA splicing, differentiation, transformation, angiogenesis, inflammation, T-cell regulation, host defense and fibrosis (*51, 52*). Based on the high expressions of stress response and inflammatory regulator, cluster 2 cells were considered as the glial and immune cells of scar. The identification of scar-associated cluster 2 serves important functions in defining the boundary of scar accurately from the cellular level.

The highest DEGs in Cluster 3 were collagen type I alpha 1 chain (Col1a1), collagen type I alpha 2 chain (Col1a2), insulin like growth factor binding protein 6 (Igfbp6) and matrix gla protein (Mgp). Col1a1 and Col1a2 belong to the type I collagen (Col I). To date, pericytes and fibroblasts have been reported to be the cell types that produce Col I after SCI (*53, 54*). Among the extracellular matrix (ECM), Col1a1 and Col1a2 were reported to be the most highly expressed in injured spinal cord of contusion SCI model at 14 dpi.

In order to further explore the function of these cell types, the Gene Ontology (GO) and Kyoto Encyclopedia of Genes and Genomes (KEGG) pathway analysis data are enriched for each cluster. The Go annotation matched well with the anatomical annotation (figure 1F). Specially, some significant GO such as synaptic vesicle cycle, regulation of neurotransmitter levels and ATP metabolic process are highly related to nomal activities of neuron in Cluster 0. Some significant GO such as myelination, axon ensheathment and positive regulation of neuron projection development are highly related to electrical signal transmission in Cluster 1. On the contrary, cluster 2 cells and cluster 3 cells exhibited damage associated patterns of gene expression, with a significant enrichment for genes that have functions in the extracellular structure organization, angiogenesis, wound healing, regulation of neuron death, regulation of inflammatory response and regeneration (figure 1J).

Based on the symmetric distribution pattern of cluster 0 cells and cluster 1 cells in normal spinal cord, we divided the T10 right lateral spinal cord into four layers (figure S1F) to further explain the relations between the subgroup and spatiotemporal distribution. After countering all samples, we found at least 1.2 mm distance from the scar center (approximately 12 spots) can cover the scar boundary. Subsequently, GSVA score of apoptosis, ferroptosis, GABAergic synapse and PI_3_K-Akt signaling pathway were respectively counted (figure S1G). We found the GSVA scores of ferroptosis and GABAergic synapse signaling pathway peaked in the scar center and were symmetrically distributed on the left and right side at three stages in Layer 3 (figures 1K-L). Layer 3 enriched by cluster 0 neurons suffered from ferroptosis after spinal cord injury, consistent with previous studies (*55*). Together, we firstly present a spatiotemporal atlas that systematically describes the spatial archetypes and cellular heterogeneity of glial scar at three stages in the early acute stage (3 dpi), subacute stage (7 dpi) and intermediate stage (14 dpi, 28 dpi) after spinal cord injury.

### Deciphering spatiotemporal distribution of resident cells in the scar

To determine the heterogeneity within the cluster 2 and 3 which represented multiple cellular components that comprise the glial scar, re-clustering analysis was performed and visualized on UMAPs (figure 2A), which revealed twelve clusters identified by annotated lineage markers (figure 2B-C). These 12 clusters represented approximately all major cell types that are known to comprise the scar including microglia, macrophages, astrocytes, oligodendrocytes, fibroblasts and endothelial cells. From the re-clustering UMAPs at the different stages after spinal cord injury (figure 2D), we found the cell spots distribution at 14 dpi and 28 dpi were similar, and the cell spots distribution at 3 dpi were significant different from that at 14 dpi and 28 dpi. By the contrast, the cell spots distribution at 7 dpi were phenotypically between cell 3 dpi spots and 14 dpi spots. The cell spots distribution provided compelling evidence about the division of time window-the early acute stage (3 dpi), subacute stage (7 dpi) and intermediate stage (14 dpi, 28 dpi) after spinal cord injury.

**Figure. 2.**
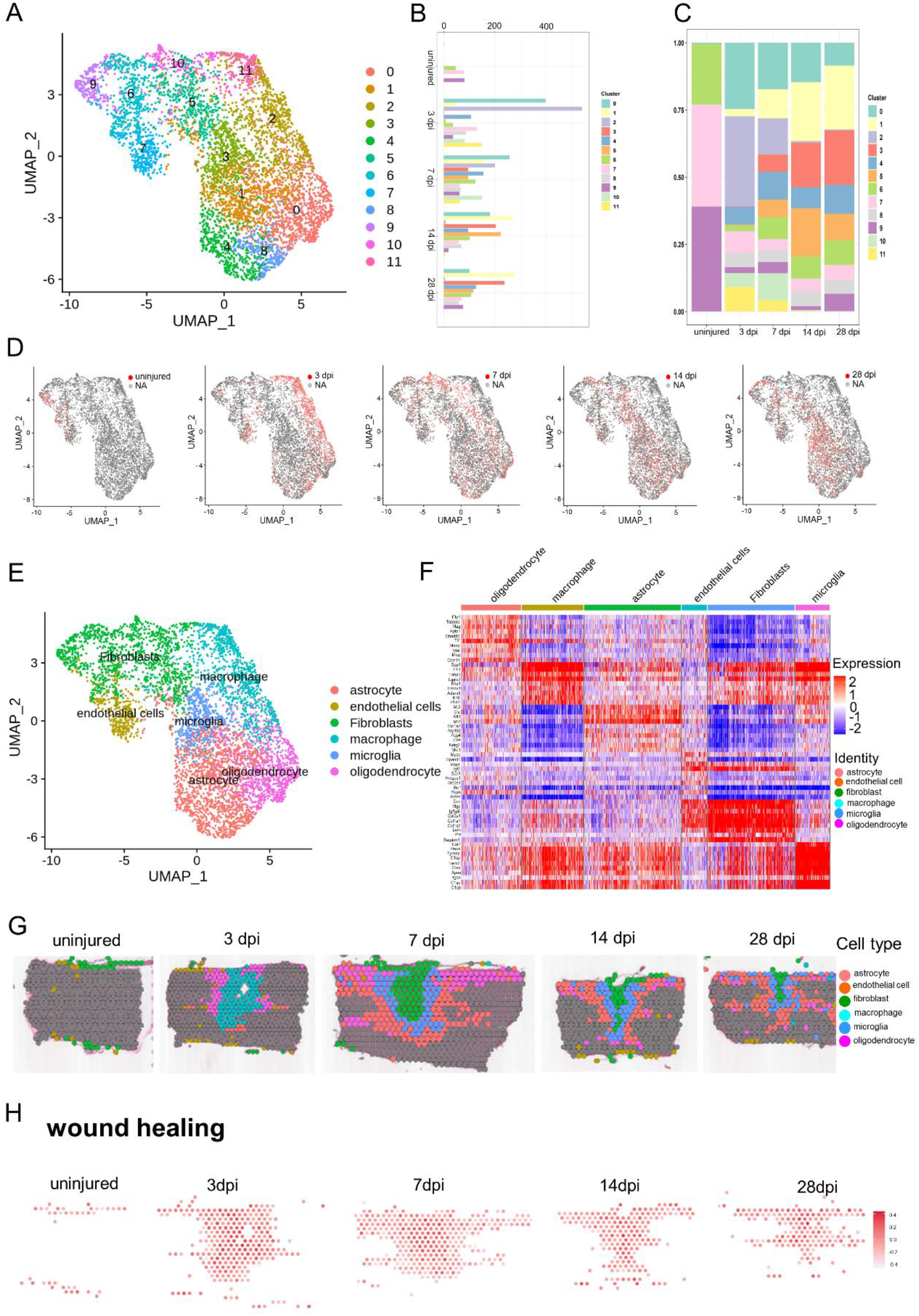
Cataloguing resident cells in the glia scar maturation with ST. (A) UMAP spots showing the re-clustering analysis of cluster 2 cells and cluster 3 cells from four stages of scar maturation. (B) Histogram showing the number of spots in each subpopulation. (C) Bar plot showing the fraction of all spots comprised by each subpopulation from different stages of scar maturation. (D) UMAP spots embedding overlay showing the distribution of spots at different time points after injury. (E) UMAP spots embedding overlay showing the six main cell clusters at different time points after injury. (F) Heat map showing the top 10 markers for individual clusters annotation shown as fraction of expression (color) of gene markers (columns). (G) The spatial maps showing the six main cell clusters at different time points after injury. (H) The spatial maps showing the GSVA score of wound healing enriched in the six main cell clusters at different time points after injury.

To gain molecular insights into the injury responses that are mediated by these cell types, we combine the subgroups and all the re-clustered celltypes into six distinct cell clusters, microglia, macrophage, astrocyte, oligodendrocyte, fibroblast and endothelial cell (figure 2E). The transcriptional signatures were shown in figure 2F. We listed the highly expressed genes in each group in detail. For example, Decorin (Dcn) was expecially expressed on fibroblast. Plp1 was expecially expressed on oligodendrocyte. Metallothionein 3 (Mt3) was expecially expressed on astrocyte. By the contrast, Spp1 was expecially expressed on multiple cellular components (such as macrophage, fibroblast and microglia). Subsequently, these six cell clusters were are mapped bact to slices (figure 2G), and we found macrophage firstly appeared in the scar area at the early acute stage (3 dpi), and then fibroblast occupied the center of the scar at the subacute stage (7 dpi). From the center of scar to the boundary, fibroblast, microglia, astrocyte and oligodendrocyte appeared in proper order at the subacute stage (7 dpi) and intermediate stage (14 dpi, 28 dpi) after spinal cord injury. We used a novel gene set analysis method (methods) infer activity maps by countering the GSVA score of cell spots in scar area. Wound healing (*56, 57*), a selected GO term, was mapped the in the scar region at three stages, and was significant active (figure 2H). Taken together, our data revealed the spatiotemporal distribution of multiple celltypes in the scar region and clearly depicted the boundary of each celltype.

### Identifying macrophage infiltrating into the scar by ST analysis

To determine the heterogeneity within the macrophage population, re-clustering analysis was performed and visualized on spatiotemporal maps (figures 3A and 3B). We identified three macrophage subtypes, which were labeled cluster 0 to 2. Cluster 0 was identified by high expression of lysozyme (Lyz2), a specific marker for macrophage, and thyrotropin releasing hormone (Trf). Cluster 0 comprised 45.3% of macrophage present at 3dpi (figures 3C and 3D). Cluster 1 expressed higher levels of platelet factor 4 (Pf4) (a specific marker for macrophage) and the anti-inflammatory marker Arginase 1 (Arg1) than Cluster 0 and 2 (figure 3E, figure S2A), comprised 51.8% of macrophage present at 3dpi. Cluster 2 expressed moderate level of Lyz2 and low level of Pf4, and comprised 2.9% of macrophage present at 3dpi (figure 3D, figure S2A).

**Figure. 3.**
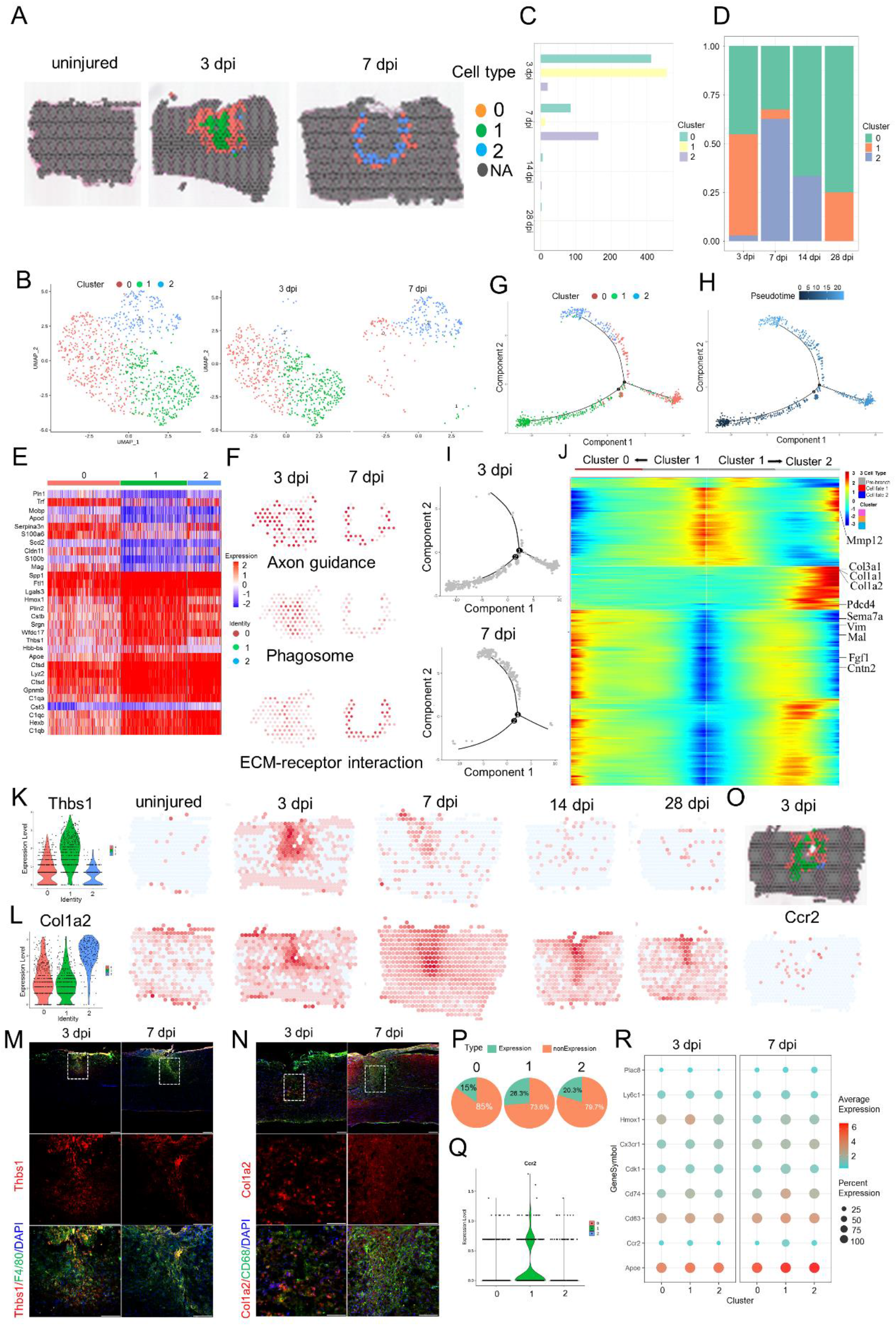
Phenotypic and functional heterogeneity of macrophage in the glia scar. (A) The spatial maps showing the distribution of three macrophage clusters in the glia scar at 3 dpi and 7 dpi. (B) UMAP spots showing the three clusters of macrophages in the glia scar at 3 dpi and 7 dpi. (C) Histogram showing the number of spots in each subpopulation at different time points after injury. (D) Bar plot showing the fraction of all spots comprised by each subpopulation at different time points after injury. (E) Heat map showing the top 10 markers for individual clusters annotation shown as fraction of expression (color) of gene markers (columns). (F) The spatial maps showing the GSVA score of axon guidance pathway enriched in cluster 0 cells, phagosome pathway enriched in cluster 1 cells and ECM-receptor interaction pathway enriched in cluster 2 cells at different time points after injury. (G, H and I) Trajectory of macrophage in the glia scar from cluster 1 cells into cluster 0/2. (J) Heat map showing the upregulated or downregulated genes in different cell fate. (K) Violin Plots and the spatial maps showing the expression of Thbs1 at different time points after injury. (L) Violin Plots and the spatial maps showing the expression of Col1a2 at different time points after injury. (M) Immunofluorescence (IF) stained 3 dpi and 7 dpi scars for Thbs1 (red), F4/80 (green) and DAPI (blue) confirming Thbs1 overexpressed in three macrophage clusters (n=3). Scare bar, 200 µm. (N) IF stained 3 dpi and 7 dpi scars for Col1a2 (red), CD68 (green) and DAPI (blue) confirming Col1a2 overexpressed in three macrophage clusters (n=3). Scare bar, 200 µm. (O) The spatial maps showing the distribution of three macrophage clusters in the glia scar at 3 dpi and the expression of Ccr2 at 3 dpi. (P) Pie chart showing the percentage of Ccr2^+^ cells in three macrophage clusters (n=27), respectively. (Q) Violin Plots showing the expression of Ccr2 in three macrophage clusters (n=27). (R) Dot plots showing the marker genes that best identifies each cell type.

Although these three subtypes did not precisely correspond to the M1/M2 division of macrophage subsets, Cluster 1 cells expressed higher levels of the Arg1 and CD14 than Cluster 0 and 2 (figure S2A). It indicates Cluster 1 cells were likely to be M2 macrophages. Interesting, ST maps show the macrophages of Cluster 1 cells were always in the central party of injury site and surrounded by the macrophages of Cluster 0. The macrophages of Cluster 2 were always in the periphery of injury site at 3 dpi (figure 3A). Subsequently, cluster 1 cells decreased dramatically and cluster 2 cells significantly increased. Interesting, cluster 2 cells interlaced with cluster 0 cells and formed a circular band of cells at 7 dpi. These three clusters cell become rare at intermediate stage (14 dpi, 28 dpi) after spinal cord injury. In addition, ST maps show the expression of macrophage marker Pf4 was significantly upregulated at 3 dpi and 7 dpi, and then back to baseline level at intermediate stage (figure S2A). There was an obvious shift in the expression of M2 macrophage marker Arg1, which peaked at 3 dpi and then fell rapidly. This was consistent with the change of macrophage phenotype after injury (*58*). The Pf4^+^ cells resided in the scar at intermediate stage (14 dpi, 28 dpi) mainly represent M1 macrophages, which resulted in an unfriend regenerate microenvironment in scar and were also characterized by CD86 and CD32 maps (figure S2A). This is the first identification of macrophage subsets spatiotemporal distribution in the scar.

It is traditionally believed that M2 macrophages could promote CNS repair while limiting secondary inflammatory-mediated injury (*58*). We found the macrophages of cluster 1 cells not only decreased dramatically, but also surrounded by the macrophages of cluster 0 and cluster 2. To investigate the functions of these clusters, we listed top 10 marker genes of these three clusters (figure 3E), and enriched the GO terms base on highly expressed genes (figure S2B). The top terms showed cluster 1 cells indeed have high enrichment factor in wound healing. However, cluster 1 cells also exhibited unique patterns of gene expression, with significant enrichments for chemotaxis, regulation of inflammatory response, response to interferon-gamma and cellular response to interleukin-1. Interesting, cluster 0 cells were associated with border and exhibited significant enrichments for axonogenesis, axon guidance, axon extension and axon ensheathment. It seems they can promote axon regeneration into the scar from both sides. The impressive functions matched the unique spatiotemporal distributions of cluster 0 cells resided in the scar periphery, which allowed cluster 0 cells to have better access to axon terminal. The GSVA score of axon guidance pathway analysis also supported this view (figure 3F). In addition, the top 4 GO terms exhibited by cluster 2 cells were extracellular structure organization, extracellular matrix organization, positive regulation of cell migration and angiogenesis. Particularly, the GO term of negative regulation of immune system process matched the traditional M2 macrophages functions limiting secondary inflammatory-mediated injury (*58*). So, we hypothesized that cluster 2 cells represent the continuators of the cluster 1 cells.

To test the hypothesis, we conducted the trajectory analysis of these three subtypes (figure 3G-J, figure S2C). Two branches were builded (figure 3G) and then the pseudo-time trajectory algorithm was applied (figure 3H). We found the evolation trajectory and transcriptional connections among the subtypes. The results indicated trajectory direction started from cluster 1 cells to a separate branch consisting of cluster 0 cells and cluster 2 cells (figure 3H-I). We observed the cell fate 1 branch enriched for genes associated with axon ensheathment, such as Sema7a, Vim, Mal, Cntn2 and Fgf1. While the cell fate 2 branch showed increased expression of extracellular matrix organization and negative regulation of immune system process related genes, such as Gpnmb, Mmp12, Col3a1, Col1a1 and Col1a2 (figure 3J).

The differentially expressed genes (DEGs) provided a molecular signature for each cell type, which were different from canonical markers (Table S3). For example, the highest DEGs in Cluster 1 were non-canonical genes such as Thbs1 (Thrombospondin 1) and Plin2 (Perilipin 2). Microvascular dysfunction is a critical pathology that underlies the evolution of secondary injury mechanisms after traumatic SCI. Thbs1 signaling was implicated in the acute neuropathological events that occur in spinal cord microvascular ECs (smvECs) following SCI (*59*). Additionally, unsatisfied intrinsic neurite growth capacity results in significant obstacles for injured spinal cord repair. Thbs1 was reported to be a neurite outgrowth-promoting molecule. Overexpression of Thbs1 in bone marrow mesenchymal stem cells (BMSCs) can promote neurite outgrowth in vitro and in vivo of Oxygen-Glucose Deprivation (OGD) treated motor neurons and SCI rat models. Specially, the transplantation of BMSCs^+^ Thbs1 could promote neurite outgrowth and functional recovery after SCI partly through the TGF-β1/p-Samd2 pathway(*60*). ST maps show the expression of Thbs1 was significantly upregulated at 3 dpi and 7 dpi, and then back to baseline level (figure 3K, figure S2C). Immunostaining showed the majority of Thbs1^+^ cells are P4/60 (a macrophage marker) positive cells at 3 dpi and 7 dpi (figure 3M). Although Spp1 was highest DEGs in Cluster 1, Spp1 expressed in multiple cell types and was unfit for non-canonical macrophage markers (figure S1E).

Another unexpected finding is that the macrophage in Cluster 2 expressed higher levels of Col I, including Col1a2, Col1a1, Col3a1, which is different from the previous study that the fibroblasts and pericytes produced Col I after SCI. Col I was highly expressed in the spinal cord, which is consistent with previous findings. Specially, Col I induced astrocytic scar formation via the integrin-N-cadherin pathway during the scar-forming phase (*61*). This was consistent with GO terms associated with extracellular matrix organization. ST maps show that the expression of Col1a2 was rare in the intact spinal cord tissues, and obviously upregulated after injury. It peaked at 7 dpi and was maintained at high level for 28 days (figure 3L). Immunostaining showed the majority of Col1a2^+^ cells are CD68 (a macrophage marker) positive cells at 3 dpi (figure 3N). Additionally, Msr1 (macrophage scavenger receptor 1) has been implicated in many macrophage-associated physiological and pathological processes including atherosclerosis, Alzheimer’s disease, and host defense. In the scar, Msr1 promote the formation of foam macrophages and neuronal apoptosis after spinal cord injury (*62*). ST maps showed that Msr1^+^ cells significantly increased at 3 dpi and 7 dpi in the injured site (figure S2D). Activation of the innate immune system promotes regenerative neurogenesis in zebrafish. TNFa from pro-regenerative macrophages induces Tnfrsf1a-mediated AP-1 activity in progenitors to increase regeneration-promoting expression of hdac1 and neurogenesis. But it is unknown whether the mammals retain the mechanism. Our ST maps showed that at least Tnfrsf1a was widespread at 3 dpi and 7 dpi in the scar (figure S2E) (*63*).

In terms of macrophage origin, the traditional view believed that bone marrow-derived mononuclear macrophages (also known as recruited macrophages) are recruited into damage tissue via chemokine (C-C motif) receptor 2 (Ccr2) in response to chemokines and inflammatory signals and affect wound healing by secreting inflammatory factors such as TNFα (*58*) (*64*). Ccr2 encodes a receptor for monocyte chemoattractant protein-1, a chemokine that specifically mediates monocyte chemotaxis, and is expressed on monocytes and macrophages, but not in microglia or tissue resident macrophage at the resting state (*65*). Recent study found the mouse meninges hosted a rich repertoire of immune cells mediating CNS immune surveillance and supplied not from the blood but by adjacent skull and vertebral bone marrow. Under spinal cord injury, local bone marrow–derived monocytes can infiltrate spinal cord from the dural meninges (*66*). We want to distinguish position and proportion of the macrophages from the blood and tissue-resident macrophages from adjacent the dural meninges. ST maps showed that Ccr2^+^ monocyte-derived macrophages were present in the scar, but their proportion in the injured site was low (figures 3O and 3P), consistent with previous study. Violin Plots showed that the expression of Ccr2 was low in these three clusters (figure 3Q). Last, we profiled the combination of the activated macrophage markers to distinguish these clusters, and found cluster 1 cells expressed higher levels of heme oxygenase 1 (Hmox1) at 3 dpi. In summary, our data reveals the spatiotemporal distribution of macrophage subtypes at the injury site after SCI.

### ST analysis reveals spatiotemporal changes in fibroblast subtypes

Fibroblasts are the main producers of the matrix (including ECM) and constitute the basic framework of tissues and organs (*67*). To assess the cellular heterogeneity among fibroblast at the injury site, re-clustering analysis was performed and visualized on spatiotemporal maps (figure 4A) and a UMAP (figure 4B-C). We identified three fibroblast subtypes, which were labeled cluster 0 to 2. Base on the spatiotemporal maps, we found cluster 2 mainly appeared in the scar at 7 dpi. Cluster 1 appeared in the scar at 7 dpi and locate in the center of scar. On the contrary, cluster 0 located in the meninges of spinal cord. The fibroblast number in Cluster 1 reached the peak at 14 dpi (figure 4D). Cluster 0 were identified by high expression of Prostaglandin D2 Synthase (Ptgds) and insulin like growth factor binding protein 6 (Igfbp6) (figure 4E). Ptgds preferentially expressed in brain and functions as a neuromodulator as well as a trophic factor in the central nervous system (CNS). In the chronic constriction injury (CCI) of sciatic nerve model, Ptgds was overexpressed in the lumbar spinal cord, but not in the striatum (*68*). Igfbp6, a member of the insulin-like growth factor-binding proteins family, modulates insulin-like growth factor (IGF) activity. In spinal cord, meningeal cells, interneurons in the deep part of the dorsalhorn and around the central canal, and motoneurons were Igfbp6 positive cells (*69*). Igfbp6 plays a key role in neuronal apoptosis after SCI. In acute SCI model, Igfbp6 was upregulated significantly and was co-localized with active caspase-3 and p53 in neurons. When Igfbp6 was knocked down, the protein levels of active caspase-3 and Bax as well as the number of apoptotic primary neurons were significantly decreased (*70*). ST maps showed Igfbp6 and Ptgds were high expression levels in Cluster 0 cells (figure S3A).

**Figure. 4.**
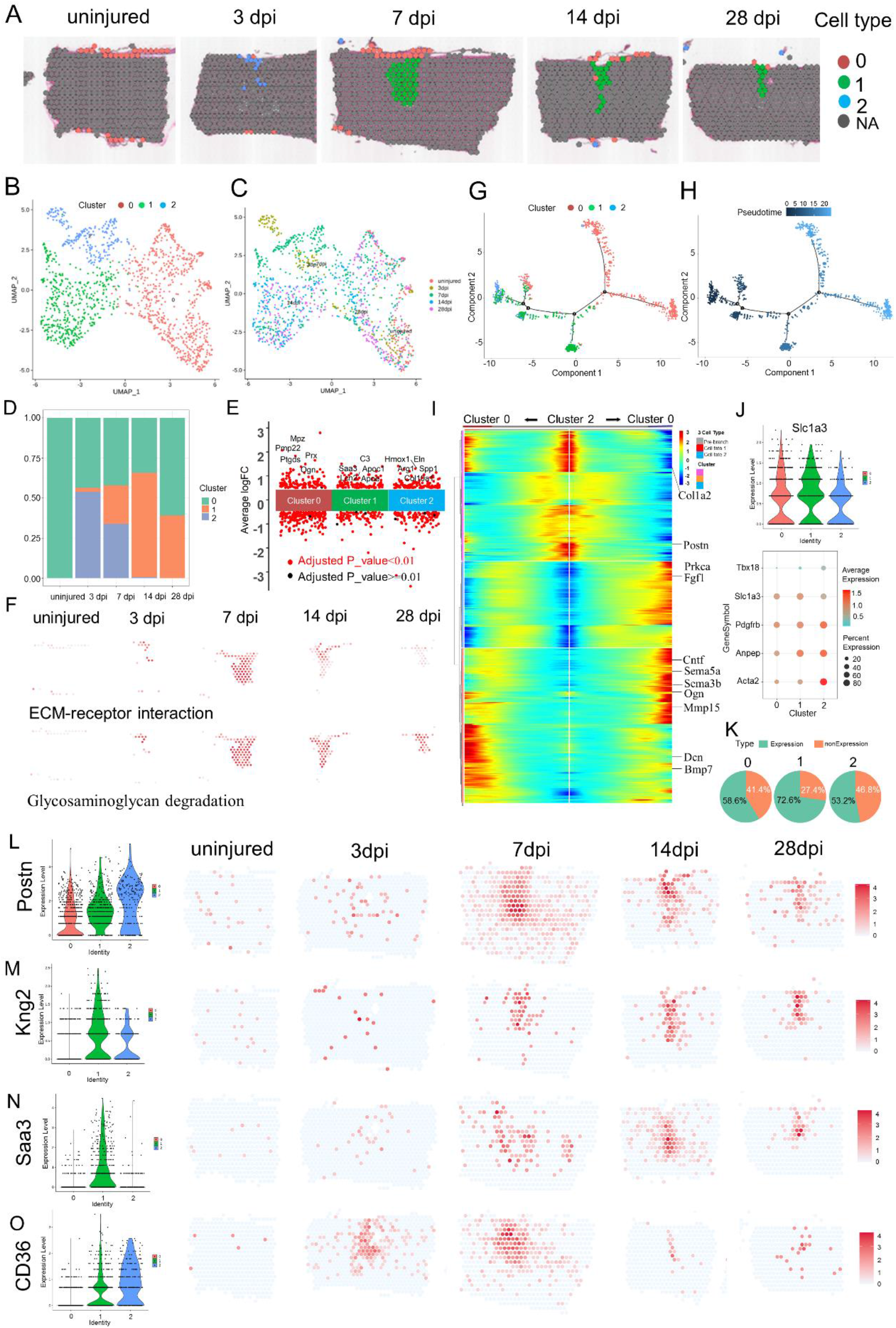
Phenotypic and functional heterogeneity of fibroblast in the glia scar. (A) The spatial maps showing the distribution of three fibroblast clusters in the glia scar at different time points after injury. (B) UMAP spots showing the three clusters of fibroblasts in the glia scar. (C) UMAP spots embedding overlay showing the distribution of spots at different time points after injury. (D) Bar plot showing the fraction of all spots comprised by each subpopulation at different time points after injury. (E) Jitterplot showing the top 5 differentially expressed genes (DEGs). (F) The spatial maps showing the GSVA score of ECM-receptor interaction pathway enriched in cluster 2 cells and Glycosaminoglycan degradation pathway enriched in cluster 1 cells at different time points after injury. (G and H) Trajectory of fibroblast in the glia scar. (I) Heat map showing the upregulated or downregulated genes in different cell fate. (J) Violin Plots showing the expression of Slc1a3 in the three clusters. (K) Dot plots showing the marker genes that best identifies type A pericytes. (L) Pie chart showing the percentage of Slc1a3^+^ cells in three fibroblast clusters (n=32), respectively. (M) Violin Plots and the spatial maps showing the expression of Postn, myofibroblast differentiation marker, at different time points after injury. (N) Violin Plots and the spatial maps showing the expression of Kng2 at different time points after injury. (O) Violin Plots and the spatial maps showing the expression of Saa3, an inflammatory ligand, at different time points after injury. (P) Violin Plots and the spatial maps showing the expression of CD36, the critical regulator of pathological skin scarring, at different time points after injury.

To investigate the functions of these clusters, we listed top 5 differential genes between these three clusters (figure 4E), and enriched the GO terms base on highly expressed genes (figure S3B). The top terms showed cluster 2 cells have high enrichment factor in wound healing. In addition, cluster 2 cells also exhibited significant enrichments for angiogenesis, extracellular matrix organization, response to transforming growth factor beta and collagen metabolic process. The GSVA score of ECM-receptor interaction pathway analysis also supported this view (figure 4F). But cluster 2 cells diminished gradually. Interesting, cluster 0 cells were associated with meninges and exhibited significant enrichments for axonogenesis, axon development, axon guidance, axon ensheathment and regulation of epithelial cell proliferation. What is more, the top 10 GO terms exhibited by cluster 1 cells were leukocyte chemotaxis, phagocytosis, regulation of immune effector process, T cell activation and activation of immune response. cluster 1 cells gradually become the main component in the scar and their enrichments were associated with inflammatory hyperactivation in the intermediate stage after spinal cord injury, consistent with previous studies (*71*). The GSVA score of glycosaminoglycan degradation in top 10 pathway analysis enriched in cluster 1 cells significantly increased from 7 dpi to 28 dpi (figure 4F), which may be related to the maturation and contracture of the scar, resulting in compartmental structures containing multiple cells (figure S3C).

To verify the evolutionary relationship between these three cell populations, we conducted the trajectory analysis (figure 4G-I). Two branches were found (figure 4G) and then the pseudo-time trajectory algorithm was applied (figure 4H). The trajectory direction started from cluster 2 cells and cluster 0 cells to a separate branch consisting of cluster 0 (figure 4H-I). We observed the cell fate 1 branch showed increased expression of extracellular matrix organization, such as Dcn, Col1a2 and Bmp7. While the cell fate 2 branch enriched for genes associated with axon guidance and axonogenesis, such as Sema3b, Sema5a, Periostin (Postn), Cntf, Ogn, Mmp15 and Fgf1 (figure 4I). Interesting, the origin of cluster 1 cells came from three directions. That is undoubtedly true that a certain percentage of cluster 1 cells came from adjacent the dural meninges after spinal cord injury. Our data showed cluster 2 cells appeared at 3 dpi and could transform into cluster 1 cells. Where these cells originate from? Previous studies reported that in mouse models that develop fibrotic tissue, the primary source of scar-forming fibroblasts was type A pericytes. Perivascular cells with a type A pericyte marker profile also exist in the human brain and spinal cord, suggesting it is conserved across diverse CNS lesions (*72, 73*). Attenuation of pericyte-derived fibrosis represents a promising therapeutic approach to facilitate recovery following spinal cord injury(*74*). We hypothesized that cluster 2 cells originate from type A pericyte and then tested the expression of type A pericyte marker GLAST (gene name Slc1a3). We verified the expression level of GLAST and GLAST^+^ cells proportion at the three clusters, and found GLAST^+^ cells were the primary origin of fibroblast in the scar (figure 4J and K), consistent with previous study(*73*).

Homeostatic fibroblasts of cluster 0 were identified based on its spatiotemporal distribution and the expression of several annotated markers of steady-state fibroblast, such as P4hb and Gsn. Homeostatic fibroblasts of cluster 0 were the predominant subtype in the uninjured spinal cord, but by 3 dpi they were replaced by the damaged-associated fibroblasts (DAF) of cluster 1 and cluster 2 (figure 4D, figure S3A). DAF were always in the center of scar. DAF were identified by high expression of Gpnmb, Ftl1, Col3a1 and Postn. Myofibroblast differentiation marker Postn is an important extracellular matrix protein and coordinate a variety of cellular processes and functions in tissue development and regeneration, including wound healing, and ventricular remodeling following myocardial infarction (*75*). Upregulation of Postn was reported in different diseases characterized by oxidative stress and inflammatory response (*76*) (*77*). In addition, upregulation of Postn suppresses SLC7A11 expression through the inhibition of p53 in VSMCs and increased the sensitivity of cells to ferroptosis. More importantly, Postn could remodel the injury environment and was identified as a therapeutic agent for traumatic injury of the CNS. Astroglial-derived Postn promotes axonal regeneration through FAK and Akt signaling pathway after spinal cord injury (*78*). But Postn was also reported to be a key player in scar formation after traumatic SCI. Genetic deletion of Postn in mice reduced scar formation at the lesion site by inhibiting the proliferation of the pericytes. Moreover, the pharmacological blockade of Postn restrained scar formation and improved the long-term functional outcome after SCI (*79*). Conversely, our ST maps found the expression of Postn predominantly in the DAF of cluster 1 and cluster 2. Although the expression of Postn also peaked at 7 dpi, it maintained a high level of expression until 28 dpi (figure 4L). We also found the non-canonical genes such as Kininogen 2 (Kng2) (figure 4M) and Serum Amyloid A3 (Saa3) displayed better specificity than canonical fibroblast markers such as P4HB which was expressed in multiple cell types (figure S3A). The expression of *Saa3* in cluster 1 increase gradually and reached the highest level at 7 dpi (figure 4N). Saa3 is a pseudogene, and acts as an inflammatory ligand and target of long non-coding RNA (lncRNA) metastasis associated lung adenocarcinoma transcript 1 (MALAT1). Saa3 was accompanied by increase of inflammatory mediators, tumour necrosis factor alpha (TNF-α) and interleukin 6 (IL-6) (*80*). Additionally, Saa3 is a key mediator of the protumorigenic properties of cancer-associated fibroblasts in pancreatic tumors (*81*). So far, the role of Saa3 in scar formation is unclear.

In spinal cord, fibrotic scar become mature gradually (figure S3C) and limits CNS regeneration in adult mammals. Similarly, skin wounds generally heal by scarring, a fibrotic process mediated by the Engrailed-1 (En1) fibroblast lineage (*82*). Preventing Engrailed-1 activation in fibroblasts yields wound regeneration without scarring. It is unknown whether fibrotic scar-forming in spinal cord need Engrailed-1 activation. In the current study, ST maps showed En1 almost was rarely expressed in spinal cord before and after injury (figure S3D), suggesting CNS having a different fibrotic scar-forming. Another study found Jun promotes hypertrophic skin scarring via CD36 in vitro and in vivo models (*83*). Indeed, we observed that injury enhance Jun expression (figure S3D). Particularly, CD36 was obviously induced in the injury and very limited to the area of scar (figure 4O). Previous study has found CD36 deletion improved recovery from spinal cord injury in mouse, but they didn’t identify the cell type expressing CD36 (*84*). In our data, there’s a fair amount of overlap between CD36^+^ spots and a-SMA^+^ (gene name Acta2) activated fibroblasts (figure S3A), suggesting CD36^+^ involved in contractures of the activated fibroblasts for scar formation. Specially, 59.83% percent of CD36^+^ cells were GLAST positive at 28 dpi. Targeting CD36 may be a direction for spinal scar therapy. Taken together, we identified the spatiotemporal distribution and origin of fibroblast subtypes at the injury site after SCI.

### ST analysis of microglia heterogeneity reveals the specific roles during gliosis

Microglia play a crucial part in scar formation (*67, 85*). To assess the cellular heterogeneity among microglia at the injury site, re-clustering analysis was performed and visualized on spatiotemporal maps (figure 5A) and a UMAP (figure S4A-B). We identified six microglia subtypes, which were labeled cluster 0 to 5 (figure 5B, figure S4C). Base on the spatiotemporal maps, we found cluster 1 cells mainly distributed in the boundary of fibroblast foci in scar at 7dpi and diminished rapidly. On the contrary, cluster 0 cells replaced cluster 1 cells in the boundary of fibroblast foci. The fibroblast number in Cluster 0 increased gradually from 7dpi to 28dpi (figure 5B). ST maps characterized the expression of P2Y12-a marker for homeostatic microglia, CD68 and Iba1-the markers for activated microglia (figure S4D).

**Figure. 5.**
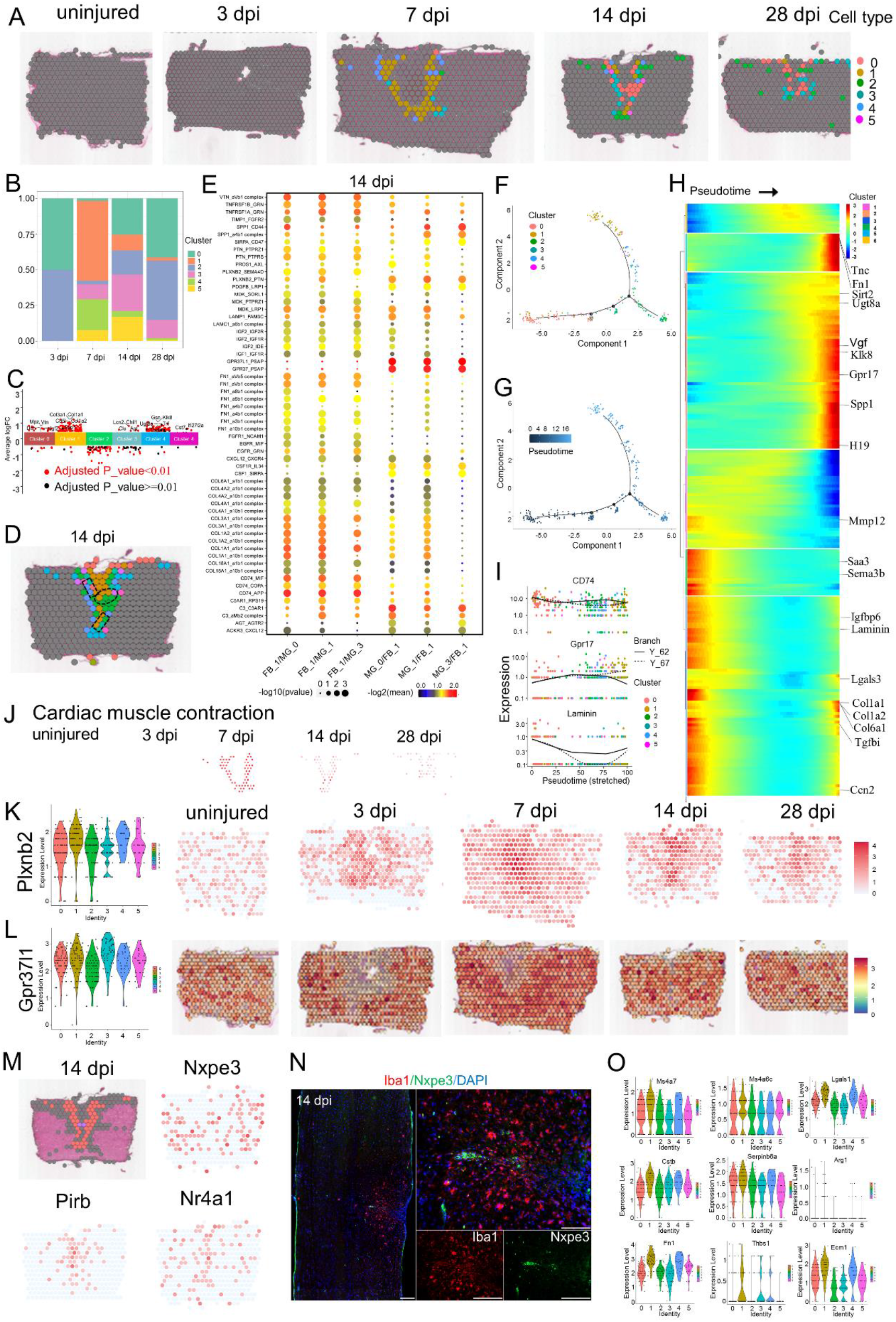
Phenotypic and functional heterogeneity of microglia in the glia scar. (A) The spatial maps showing the distribution of six microglia clusters in the glia scar at different time points after injury. (B) Bar plot showing the fraction of all spots comprised by each subpopulation at different time points after injury. (C) Jitterplot showing the top 5 differentially expressed genes (DEGs) at each subpopulation. (D) The definition of the boundary areas to study the interaction between two neighbor microglia clusters in the scar. The regions with 2 spots wide along the boundary lines in each cluster were selected. (E) Dot plot showing the mean interaction scores between the neighbor clusters at the boundaries for ligand-receptor pairs. The ligand-receptor pairs are listed on the left. Size of the circle denoted p-value, and color denoted the mean interaction scores. (F and G) Trajectory of microglia in the glia scar. (H) Heat map showing the changes in genes expression along a spatial trajectory. (I)The expression of CD74, Gpr17 and Laminin correlating with pseudotime. (J) The spatial maps showing the GSVA score of cardiac muscle contraction pathway enriched in cluster 1 cells at different time points after injury. (K) Violin Plots and the spatial maps showing the expression of Plxnb2 at different time points after injury. (L) Violin Plots and the spatial maps showing the expression of Gpr37l1 at different time points after injury. (M) The spatial maps showing the expression of selected DEGs in the spots that crossed the fibroblasts scar and formed bridges at 14 dpi. (N) IF stained 14 dpi scars for Nxpe3 (red) and DAPI (blue). Scare bar, 200 µm. (O) Violin Plots showing the expression of selected genes contributing to microglia-organized scar-free spinal cord repair in neonatal mice.

To characterize the functions of these clusters, we listed top 5 differential genes for each cluster (figure 5C), and enriched the GO terms base on highly expressed genes (figure S4E). The top terms showed cluster 0 cells have high enrichment factor in axon ensheathment, axon development and ensheathment of neurons. The pathway of ECM-receptor interaction was also significantly enriched in cluster 0 cells. Interesting, ST maps showed the minority of cluster 0 cells crossed the boundary of fibroblast scar. Cluster 1 cells, cluster 2 cells, cluster 3 cells exhibited significant enrichments for angiogenesis, extracellular matrix organization and extracellular structure organization. The GSVA score of cardiac muscle contraction pathway analysis peaked at 7 dpi and might explained the phenomenon of scar contracture (figure 5J). In addition, cluster 5 cells exhibited significant enrichments for negative regulation of cell motility, negative regulation of cell migration, negative regulation of locomotion and r negative regulation of immune system process, which were similar to the function of neonatal microglia (*85*). Unfortunately, cluster 5 cells are few, and was not enough to change the whole inflammatory environment.

We next investigated the communication and interaction between microglia and fibroblast, the interface regions of clusters were selected with the range of at least 2 spots for each cluster. The top enriched gene-pairs included TNFRSF1B_GRN, TNFRSF1A_GRN, SPP1_CD44, PLXNB2_PTN, GPR37L1_PSAP and GPR37_PSAP (figures 5D and 5E, figures S4F and S4G). For example, PLXNB2 are transmembrane receptors that participate in axon guidance and cell migration in response to semaphorins (*86*). Plexin B2 was induced in injury-activated microglia and macrophages early after SCI and facilitated the formation of concentric rings of microglia and astrocytes around the necrotic core of the lesion in a mouse model of SCI (*87*). ST maps showed Plexnb2^+^ cell enriched in scar area (figure 5K). Additionally, GPR37L1_PSAP and GPR37_PSAP were top 2 enriched gene-pairs between microglia cluster 0/1/3 and fibroblast cluster 1 from 14 dpi to 28 dpi. Prosaposin (gene name PSAP) can be secreted from various cell types in response to cellular stress. Secreted PSAP can initiate endocytose or pro-survival signaling pathways via binding to GPR37 and GPR37L1 (*88, 89*). ST maps showed PSAP was enriched in scar-resident fibroblasts (data not shown) and GPR37L1 was enriched in scar-resident microglia (figure 5L), but the exact role of fibroblast secreted PSAP s should be explored in future.

Subsequently, we conducted the trajectory analysis (figure 5F-I) and the pseudo-time trajectory algorithm (figure 5G). The trajectory direction started from cluster 1 cells and cluster 0 cells to a separate branch consisting of cluster 2-5 (figure 5H). We observed that the expression level of enriched genes associated with axon ensheathment and axon development, such as Tnc, Fn1, Sirt2, Ugt8a and Klk8, gradually increased following the scar stabilization. On the contrary, Sema3b, Mmp12 and Laminin gradually declined. We also observed that the expression level of G protein-coupled receptor 17 (Gpr17) significantly increased. Gpr17, a P2Y-like receptor, may act as a ‘sensor’ of damage that is known to be activated by both uracil nucleotides and cysteinyl-leukotriene released in the lesioned area, and could also participate in post-injury responses (*90*), In non-injured spinal cord parenchyma, Gpr17 is present on a subset of neurons and of oligodendrocytes at different stages of maturation, whereas it is not expressed by astrocytes. Induction of SCI resulted in marked cell death of Gpr17^+^ neurons and oligodendrocytes inside the lesion followed by the appearance of proliferating Gpr17^+^ microglia/macrophages migrating to and infiltrating into the lesioned area (*91*). In vivo pharmacological or biotechnological knock down of Gpr17 markedly prevents brain infarct evolution, suggesting Gpr17 as a mediator of neuronal death at this early ischemic stage (*92*). Gpr17 also acts as an intrinsic timer of oligodendrocyte differentiation and myelination. Pharmalogical targeting of Gpr17 signaling in OPCs and microglial inhibition of oligodendrocyte maturation together promote robust myelination of regenerated axons after CNS injury (*93*). The classically activated neuroinflammatory microglia induce A1 astrocytes by secreting Il-1α, TNF and C1q, and that these cytokines together are necessary and sufficient to induce A1 astrocytes (*94*). ST maps characterized the enhanced expression of C1qa in six microglia subtypes at different time point (figure S4H).

Recent study found microglia-organized scar-free spinal cord repair in neonatal mice. Fibronectin^+^ microglia in cluster 3 (MG3) formed bridges between the two stumps starting from 3 dpi. Activated microglia were observed inside the lesion in the absence of GFAP^+^ astrocytes and collagen I^+^ fibroblasts (*85*). In our adult mouse model, ST maps showed A few microglia in cluster 0 crossed the fibroblasts scar and formed bridges between the two stumps at 14 dpi. Unfortunately, the bridges eventually disappeared at 28 dpi. We profiled the selected cells that crossed the fibroblasts scar and found Nxpe3, Pirb and Nr4a1 specificity highly expressed in these cells (figure 5M). IHC verified the expression of Nxpe3 in the boundary of fibroblast scar (figure 5N). Finally, we profiled ECM-related genes (for example, Fn1, ECM1 and Thbs1), wound healing genes (for example, Anxa1), positive regulation of cell adhesion (for example, Spp1), negative regulation of hydrolase activity (for example, Cstb, Serpinb6a and Stfa1), negative regulation of immune system process (for example, Arg1), disease-associated genes (for example, Igf1 and Clec7a), embryonic stem cell-related genes (for example, Ms4a7, Lgals and Ms4a6c), phospholipase inhibitor activity (for example, Anxa2, Anxa5 and Apoc1) in six microglia subtypes. These genes were highly expressed in MG3. But, we found that Stfa1 and Clec7a weren’t expressed in six microglia subtypes, the expression level of Arg1 and Thbs1 were very low (figure 5O). Specially, Anxa1, a potent anti-inflammatory protein, may contribute to the ability of MG3 cells to mediate rapid inflammation resolution and were barely expressed in the microglia of cluster 2 and cluster 3, which were the two major types of microglia in the scar at 28 dpi (figure S4I anf figure 5B). These difference between the microglia in six clusters with the MG3 cells in neonatal mice may contribute to decipher the difference in regenerative ability between adult and neonatal mice.

### Six different astrocyte subtypes are identified by ST

Astrocytes are thought to be the main drivers of encirclement. Tightly connected astrocytes continuously reshape the boundary at the edge of injury, wrapping immune cells and fibroblast-like cells with ephrin-mediated cell adhesion and spatially isolating remaining neural tissue from injury and fibrosis (*56, 71, 95*). We next assessed the cellular heterogeneity among astrocyte at the scar, re-clustering analysis was performed and visualized on spatiotemporal maps (figure 6A) and a UMAP (figure S5A-B). We identified six astrocyte subtypes, which were labeled cluster 0 to 5 (figure 6B, figure S5C). Base on the spatiotemporal maps, we found cluster 2 cells mainly distributed in the grey matter layer of microglia concentric rings in scar at 7dpi and diminished rapidly. Cluster 2 cells might represent protoplasmic astrocytes. On the contrary, cluster 0 cells mainly distributed in the white matter layer of scar, suggesting these cells represented fibrous astrocyte. The astrocyte number in Cluster 0 peaked at 14 dpi (figure 6B). The astrocyte number in cluster 1 continuously increase. ST maps characterized the expression of GFAP and Lcn2-the markers for activated astrocyte (figure S5D-E).

**Figure. 6.**
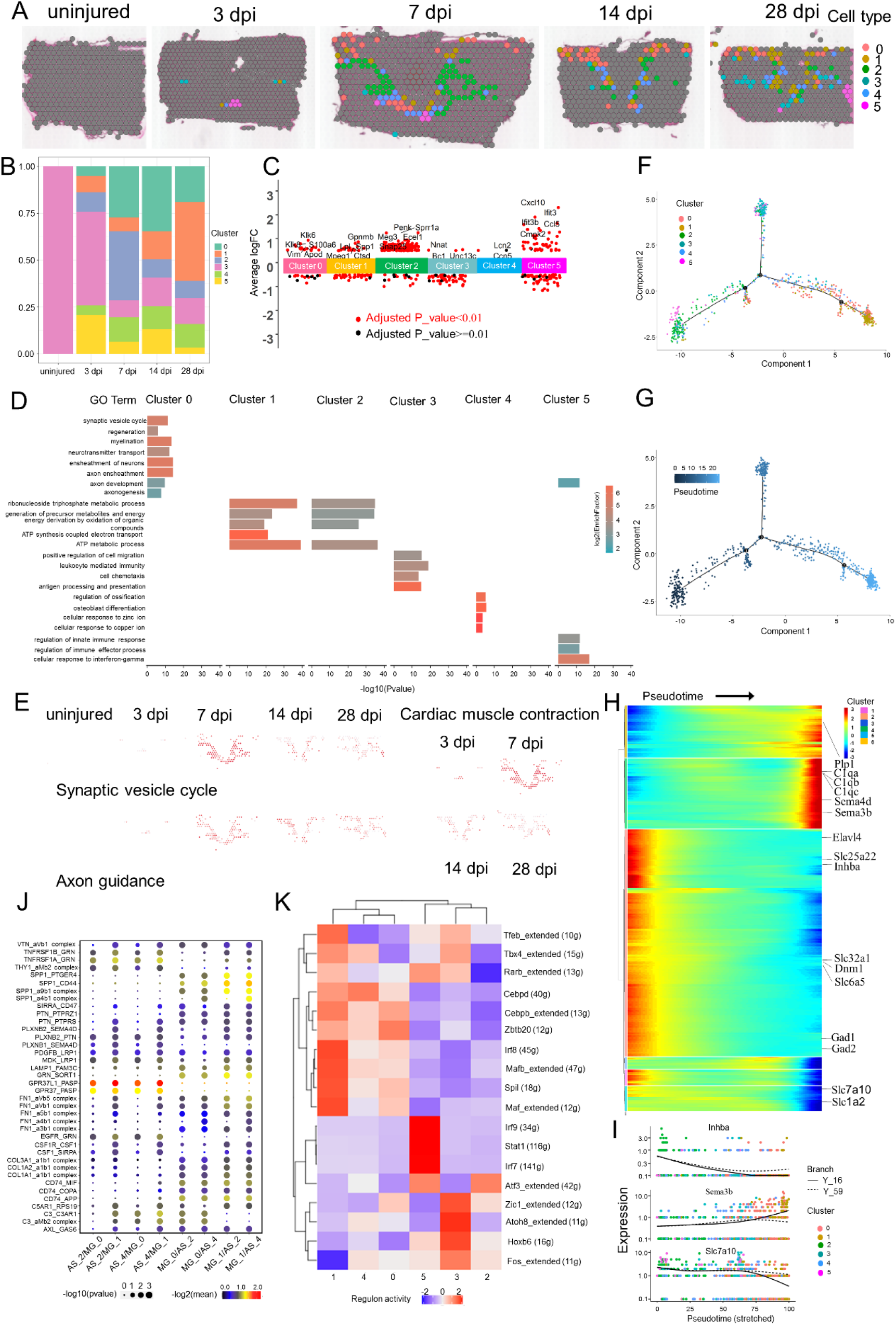
Phenotypic and functional heterogeneity of astrocyte in the glia scar. (A) The spatial maps showing the distribution of six astrocyte clusters in the glia scar at different time points after injury. (B) Bar plot showing the fraction of all spots comprised by each subpopulation at different time points after injury. (C) Jitterplot showing the top 5 differentially expressed genes (DEGs) at each subpopulation. (D) Selected GO terms that were enriched for each cluster. Fish’s exact test in the pathway enrichment was used. (F) The spatial maps showing the GSVA score of synaptic vesicle cycle pathway enriched in cluster 0 cells, axon guidance pathway enriched in cluster 0 cells and cardiac muscle contraction pathway enriched in cluster 2 cells at different time points after injury. (F and G) Trajectory of astrocyte in the glia scar. (H) Heat map showing the changes in genes expression along a spatial trajectory. (I)The expression of Inhba, Sema3b and Slc7a10 correlating with pseudotime. (J) Dot plot showing the mean interaction scores between the neighbor clusters at the boundaries for ligand-receptor pairs. The ligand-receptor pairs are listed on the left. Size of the circle denoted p-value, and color denoted the mean interaction scores. (K) Heat map showing the enriched transcription factors by single-cell regulatory network inference and clustering (SCENIC) analysis at different clusters after injury.

To further explain the biological process of the subgroup, we characterized top 5 differential genes for each cluster (figure 6C), and enriched the GO terms base on highly expressed genes (figure 6D). The top terms showed cluster 0 cells have high enrichment factor in traditional functions (*96*), such as neurotransmitter transport and synaptic vesicle cycle. In addition, cluster 0 cells also exhibited significant enrichments for myelination, axonogenesis, axon ensheathment, axon development and regeneration. The GSVA score of synaptic vesicle cycle pathway analysis and axon guidance also supported this view (figure 6E). Cluster 1 cells also exhibited significant enrichment for ATP metabolic process. The corresponding KEGG analysis showed cluster 1 cells have high enrichment factor in cardiac muscle contraction pathway. The GSVA score showed it peaked at 7dpi around the microglia scar (figure 6E). The top terms showed cluster 2 cells have high enrichment factor in regulate brain energy metabolism and homeostasis (*56*), such as ATP metabolic process and generation of precursor metabolites and energy. Interesting, cluster 4 cells exhibited significant enrichments for osteoblast differentiation, regulation of ossification, cellular response to zinc ion and cellular response to copper ion. Astrocytes were known to remove excessive potassium ions in the microenvironment (*97*). Here, we found cluster 4 cells contributed to the homeostasis in zinc ion /copper ion, which caused neuronal cell death (*98*).

We next conducted the trajectory analysis (figure 6F) and the pseudo-time trajectory algorithm (figure 6G). The trajectory direction started from cluster 2 cells to a separate branch consisting of cluster 0, 1, 3, 4, 5. We observed the genes with significantly elevated levels of expression were associated with immune response Lectin induced complement pathway, such as complement C1q Chain A (C1qa), complement C1q Chain B (C1qb) and complement C1q Chain C (C1qc) (figure 6H). Slute carrier family 25 member 22 (Slc25a22), solute carrier family 7 member 10 (Slc7a10) and solute carrier family 1 member 2 (Slc1a2) were involved in homeostasis of excitatory neurotransmitter glutamate, and markedly decreased. At the same time, solute carrier family 32 member 1 (Slc32a1), involved in gamma-aminobutyric acid (GABA) and glycine uptake into synaptic vesicles, also markedly declined. The relation between the local excitatory disorder after SCI and these transporters downregulation remain unknow (*41*). In addition, the pro-regeneration molecular inhibin subunit beta A (Inhba) markedly declined followed by the increase of Semaphorin 3B (Sema3b) (figure 6I), which inhibits axonal extension. These changes suggested the microenvironment of the astrocyte scar switched to inhibit regeneration with the maturation of scar (figure S5F) (*57, 99*).

Subsequently, we next investigated the communication between astrocyte and microglia. The top enriched gene-pairs included TNFRSF1B_GRN, TNFRSF1A_GRN, SPP1_PTGER4, SPP1_CD44, SPP1_a9b1 complex, SPP1_a4b1 complex, PLXNB2_SEMA4D, PLXNB2_PTN, GPR37L1_PSAP, GPR37_PSAP, CD74_MIF and CD74_APP (figure 6J). Interesting, GPR37L1_PSAP, GPR37_PSAP and PLXNB2_SEMA4D were in the top relationship between microglia and fibroblast. Here, these pairs were still critical. For example, GPR37L1_PSAP and GPR37_PSAP were top 2 enriched gene-pairs between astrocyte cluster 2/4 and microglia cluster 0/1. Additionally, the expression level of Sema4d gradually increased, and the pair of PLXNB2_SEMA4D between astrocyte cluster 2/4 and microglia cluster 1 has been enriched. These data indicated there were some common mechanisms mediate cell connections in the scar.

Finally, we profiled the single-cell regulatory network inference and clustering (SCENIC) analysis(*100*) and found the six subgroups were grouped into two major groups (figure 6J). The cell number of groups 1 (cluster 0, cluster 1 and cluster 4) increased gradually. Groups 1 cells might represent the traditional A1 astrocyte (*94*). In addition, ST maps showed that group 1 cell wrapped microglial scar at intermediate stage (14 dpi, 28 dpi). Interesting, we found 10 transcription factors drived astrocyte transformation to the A1 like-groups 1 cells. Additionally, Eph/ephrin signaling can the activated astrocytes maintain the homeostasis of extracellular glutamate. The EphA4 signaling prevents glutamate excitotoxicity (*101*). ST maps showed that the expression level of EphA4 was quite low, suggesting the neurotransmitter disorder in scar (figure S5G). Transglutaminase 2 (TG2) plays a key role in regulating the response of astrocytes to damage. Astrocytic-specific TG2 deletion or inhibition results in a significant improvement in functional recovery after SCI (*102*). ST maps showed TG2 was upregulated in astrocytes in scar at different time point after SCI (figure S5H). Glial BAI3 and BAI1 binding to RTN4/NoGo receptors regulate axonal elongation and synapse formation, thereby controlling neural network activity (*103*). Our ST showed the expression level of Bai3 in astrocyte scar was low at different time point after SCI (figure S5I). Additionally, astrocyte-derived saturated lipids contained in APOE and APOJ lipoparticles mediate toxicity. Eliminating the formation of long-chain saturated lipids by astrocyte-specific knockout of the saturated lipid synthesis enzyme ELOVL1 mitigates astrocyte-mediated toxicity (*104*). Our ST showed the activated astrocytes in scar have high expression of Apoe, Apoj and Elovl1 (figure S5J). An unexpected finding was that the activated astrocytes in scar have high expression of Clu (figure S5K), an anti-neuroinflammatory gene. The previous study showed that CLU was secreted by Schwann cells in peripheral sciatic nerve and induced outgrowth of sensory neurons, but not motor neurons (*50*). Recently, an exciting result showed ‘runner plasma’, collected from voluntarily running mice and infused into sedentary mice, reduces baseline neuroinflammatory gene expression and experimentally induced brain inflammation. Mechanically, the complement cascade inhibitor CLU in plasma reduced neuroinflammatory gene expression in a mouse model of acute brain inflammation and a mouse model of Alzheimer’s disease (*105*). The exact role of Clu in spinal cord injury remains unknown. In a word, these findings call for a reinterpretation of astrocyte infiltration into the scar and may inform future therapeutic approaches.

### ST identified the different oligodendrocyte subtypes in the scar

Oligodendrocyte wrapped around the axons of neurons in the CNS, forming myelin insulators on the surface of neurons that allow electrical signals to travel more efficiently. Interesting, subpopulations of oligodendrocyte and their progenitor cells have much in common with immune cells in mouse models of multiple sclerosis, and they can be involved in clearing myelin sheaths damaged by disease, in a manner similar to immune cells (*106*). We next assessed the cellular heterogeneity among astrocyte at the scar, re-clustering analysis was performed and visualized on spatiotemporal maps (figure 7A) and a UMAP (figure S6A-B). We identified three oligodendrocyte subtypes, which were labeled cluster 0 to 2 (figure 7B, figure S6C). Base on the spatiotemporal maps, we found cluster 0 cells mainly distributed in the outer layer of astrocyte concentric rings in scar at 3 dpi and diminished rapidly. On the contrary, cluster 2 cells mainly distributed in the white matter of scar (figure 7A). The oligodendrocyte number in cluster 1 continuously increase. ST maps characterized the expression of Mbp, Mog, Mag, and Cldn11-the markers for activated astrocyte (figure S6D-E).

**Figure. 7.**
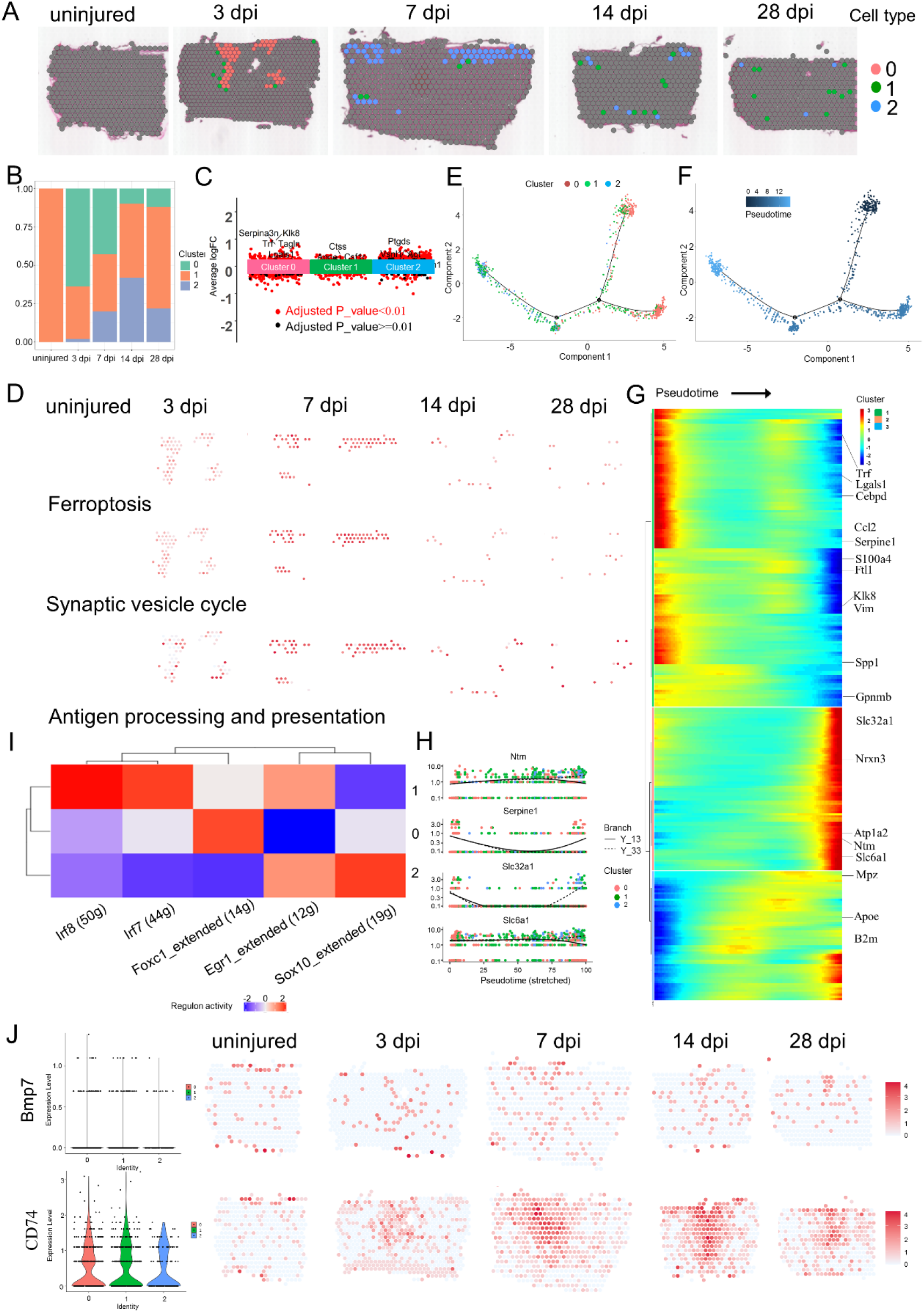
Phenotypic and functional heterogeneity of oligodendrocyte in the glia scar. (A) The spatial maps showing the distribution of three oligodendrocyte clusters in the glia scar at different time points after injury. (B) Bar plot showing the fraction of all spots comprised by each subpopulation at different time points after injury. (C) Jitterplot showing the top 5 differentially expressed genes (DEGs) at each subpopulation. (D) The spatial maps showing the GSVA score of ferroptosis pathway enriched in cluster 0 cells, antigen processing and presentation pathway enriched in cluster 1 cells and synaptic vesicle cycle pathway enriched in cluster 2 cells at different time points after injury. (E and F) Trajectory of oligodendrocyte in the glia scar. (G) Heat map showing the changes in genes expression along a spatial trajectory. (H)The expression of Ntm, Serpine1, Slc32a1 and Slc6a1 correlating with pseudotime. (I) Heat map showing the enriched transcription factors by single-cell regulatory network inference and clustering (SCENIC) analysis at different clusters after injury. (J) Violin Plots and the spatial maps showing the expression of Bmp7 at different time points after injury. (K) Violin Plots and the spatial maps showing the expression of CD74 at different time points after injury.

To further explain the biological process of the subgroup, we characterized top 5 differential genes for each cluster (figure 7C), and enriched the GO terms base on highly expressed genes (figure S6F). The top terms showed cluster 0 cells have high enrichment factor in response to wounding. In addition, cluster 0 cells also exhibited significant enrichments for epithelial cell proliferation, regeneration, angiogenesis, axon regeneration and glial cell migration. The GSVA score of p53 signaling pathway analysis and also supported this view (figure 7D). But cluster 0 cells diminished gradually. Additionally, cluster 1 cells exhibited significant enrichments for neutrophil activation involved in immune response, leukocyte mediated immunity, antigen processing and presentation and macrophage activation. The GSVA score of antigen processing and presentation analysis enriched in cluster 1 cells significantly increased from 7 dpi to 28 dpi (figure 7D), suggesting they can work in a manner similar to immune cells (*106*). Cluster 2 cells exhibited significant enrichments for neurotransmitter transport, regulation of synaptic plasticity and neurotransmitter secretion. The GSVA score of GABAergic synapse in top 3 pathway analysis enriched in cluster 2 cells significantly increased from 7 dpi to 28 dpi (figure 7D), which may be related to the inhibition of neuronal excitability after injury (*41*).

We next conducted the trajectory analysis (figure 7E) and the pseudo-time trajectory algorithm (figure 7F). The trajectory direction started from cluster 0 cells to a branch consisting of cluster 1 and 2. We observed the expression level of serpin family E member 1 (Serpine1), a component of innate immunity, gradually declined. But beta-2-microglobulin (B2m), a component of the class I major histocompatibility complex (MHC), was involved in the presentation of peptide antigens to the immune system and gradually increased. Galectin 1 (Lgals1), playing a role in regulating apoptosis, gradually declined. Interesting, solute carrier family 6 member 1 (Slc6a1) and solute carrier family 32 member 1 (Slc32a1) mediated the rapid removal of GABA and maintain low extracellular levels. The expression level of them significantly increased, suggesting oligodendrocyte also participated in the regulation of excitability in the scar. In addition, cell adhesion molecules neurotrimin (Ntm) exerts an inhibitory impact on neurite outgrowth (*107*), whereas its expression level significantly increased during the maturation of scar.

Finally, we profiled SCENIC analysis (figure 7I) and found Foxc1 potentially controlled cluster 0 cells. Foxc1 was as an important regulator of cell viability and resistance to oxidative stress in the eye (*108*). Irf7 and Ifr8 controlled cell fate of cluster 1 cells. Interesting, Sox10 controlled cell fate of cluster 2 cells. Sox10 were verified to play a central role in oligodendrocyte maturation and CNS myelination. Specifically, Sox10 activates expression of myelin genes, during oligodendrocyte maturation, such as Dusp15 and Myrf (*109*).

Oligodendrocyte death after SCI contributes to demyelination of spared axons. Bone morphogenic protein-7 (BMP7) could potently inhibit TNF-α-induced oligodendrocyte apoptosis (*110*). Overexpression of BMP7 reduced oligodendrocytes loss and promoted functional recovery after spinal cord injury (*111*). In our data, ST maps showed cells secreted BMP7 were mainly in the fibroblast scar, suggesting its beneficial role (figure 7J). Last, ST maps characterized the expression of CD74-a component of the class II major histocompatibility complex (MHC) (figure 7K). CD74 is a critical chaperone that regulates antigen presentation for immune response (*112*). All the three oligodendrocyte subtypes have a moderate expression of CD74. Taken together, these findings call for a reinterpretation of oligodendrocyte infiltration into the scar and deciphered the potentially functions.

### Gene regulatory networks controlling the scar programming

To better understand scar-relevant changes in genes regulation and interactions between cell types, we carried out WGCNA and an unbiased coexpression analysis (*38*) of our ST data. In WGCNA, 2000 genes (Table S4) from the spatiotemporal transcriptome dataset in cluster 2 and cluster 3 (figure 1H) are clustered. The branches of highly correlating genes were formed, which were cut and assigned a color. Finally, the correlation of genes and modules was calculated (figure 8A), and 14 modules were identified (figure 8B and figure S7A). Among the 14 modules, the MEgreenyellow module had the most significant correlation with the time (figure S7A), which gradually increasing following the scar formation and approximately representing microglia and astrocyte (figure 8B). We next conducted the network of the enriched hub module genes involved in Megreenyellow to identify key controlling genes (figure 8C). Meblue module and Meblack module had the most significant correlations with the early acute stage (3 dpi) and represnting macrophage and oligodendrocyte, and hub module genes in Meblue (figure 8D) and in Meblack (figure S7B) were respectively carried out.

**Figure. 8.**
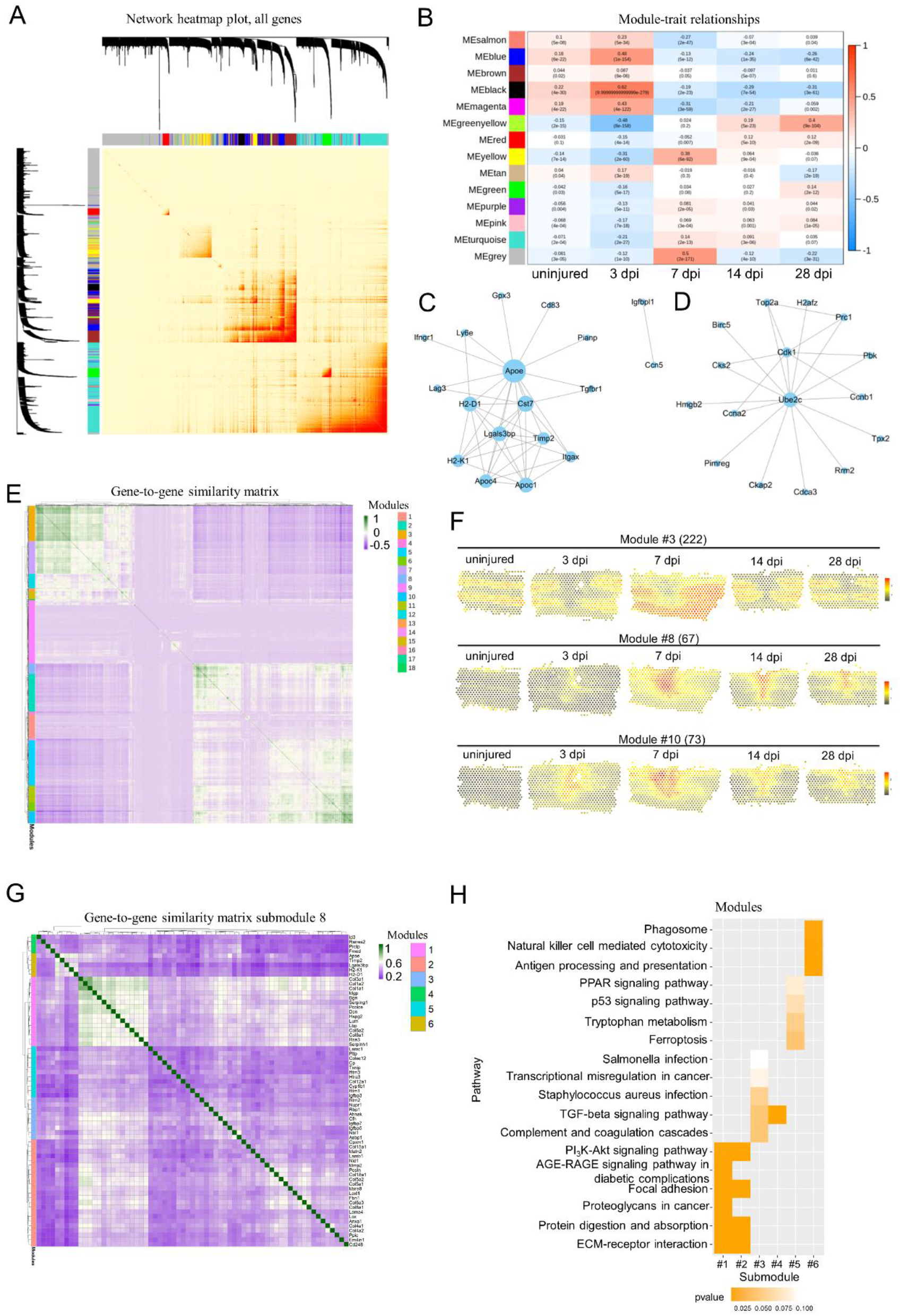
Spatiotemporal dynamics of genes expression during the maturation of scar. (A) Hierarchical clustering showing the network about the correlation of module membership and gene significance, as well as the correlation among all genes. (B) The heatmap depicting the correlation scores (digit in the box above) as well as its corresponding P-value (digit in the box below) of modules (rows) and different time after injury (columns). (C) Co-expression of selected hub genes enrichment in Megreenyellow. The networks were created by Cytoscape. (D) Co-expression network of selected hub genes enrichment in Meblue. The networks were created by Cytoscape. (E) Biclustering of the mouse spatial gene expression measurements revealing spatially and temporally coexpressed genes. Identifiers of coexpression modules are listed. (F) Average spatiotemporal expression dynamics of modules 3, 8 and 10. (G) Hierarchical clustering of gene in module 8. (H) Analysis of enriched KEGG pathway among the genes for submodules in (G).

Subsequently, we adapted another unbiased coexpression analysis more suitable for ST data (*38*). We identified 18 major coexpression modules on the basis of 2,000 highly variable genes from all spatiotemporal transcriptome dataset (Table S5). Among them, module 3, 8, 10, 16, and 17 are highly correlated and used as candidate analysis modules (figure 8E). ST maps showed the spatiotemporal activities of module 3, 8 and 10 (figure 8F). We next grouped the genes of module 3, 8 and 10 on the basis of their expression pattern, resulting in submodules (figure 8G, figure S7C-7F). Col3a1, col1a2 and col1a1 (submodule 8.1) have highest corelation score (figure 8G). KEEG pathways enrichment analysis of the submodules 8.1 show that they are enriched for ECM-receptor interaction (figure 8H). Gpnmb, Lyz2, Ftf1, Lgals3, Ctsd and Ctsb (submodule 10.3) have highest corelation score (figure S7E). KEEG pathways enrichment analysis of the submodule 10.3 show that they are enriched for lysosome (figure S7F). Collectively, the pathway activities encompassed by module 8 and 10 reveal signaling within and between cell types during glial and fibroblastic activation in the scar.

### The scar boundary was reinterpreted by ST

The traditional characteristics of lesion site in mice after adult injury were scare structure visualization with immunofluorescence for laminin, fibronectin, chondroitin sulfate proteoglycan (CSPG), GFAP, CD68, CD31 and collagen (*85*). All these data came from the sections and we lack a global understanding of its distribution throughout the spinal cord scar. Although diffusion tensor imaging (DTI) could accurately provide information of cavity area after SCI, this method is difficult to distinguish cell types (*113, 114*). With the help of tissue transparency, laminin, fibronectin and collagen were re-characterised in our mice model of T10 right lateral hemisection (figure 9A). Laminin was highly enriched in the wound area at 7 dpi. On the contrary, fibronectin and collagen was relatively low expression, which were different form the expression pattern of neonatal mice in spinal cord crush. We next applied our ST data to exhibit the laminin (figure 9B), fibronectin (figure 9C). Overall, the ST maps showed fibronectin was upregulated in the injury site and roughly characterised the region of scar in the early acute stage (3 dpi), subacute stage (7 dpi) and intermediate stage (14 dpi). On the contrary, the ST maps showed laminin was upregulated in the injury site and clearly characterised the region of scar in the subacute stage and intermediate stage. The ST maps of CD31 showed the neovascularization failed to penetrate the scar (figure 9D), which was similar to slices staining (*85*). Collectively, it is a potential breakthrough that combining the ST maps of scar markers to identify the size of scar. To define the distribution of each cell type more accurately, we counted the number and fraction of each cell type maintaining scar architecture at different time after SCI (figure 9E and 9F), and found that fibroblasts are always in the center during scar shrinkage and maturation. Fibrotic scars are surrounded by microglial scars, which are wrapped by the astrocytic scars (*115*). Specially, microglial scars and astrocytic scars did shrink at the gray matter. But they expanded a larger area along white matter at 28 dpi (figure 9E and 9F). Taken together, we identified the spatiotemporal dynamics of key genes in specific cell types, thus providing a global atlas of gene expression and molecular regulation driving scar formation following SCI (figure 9G).

**Figure. 9.**
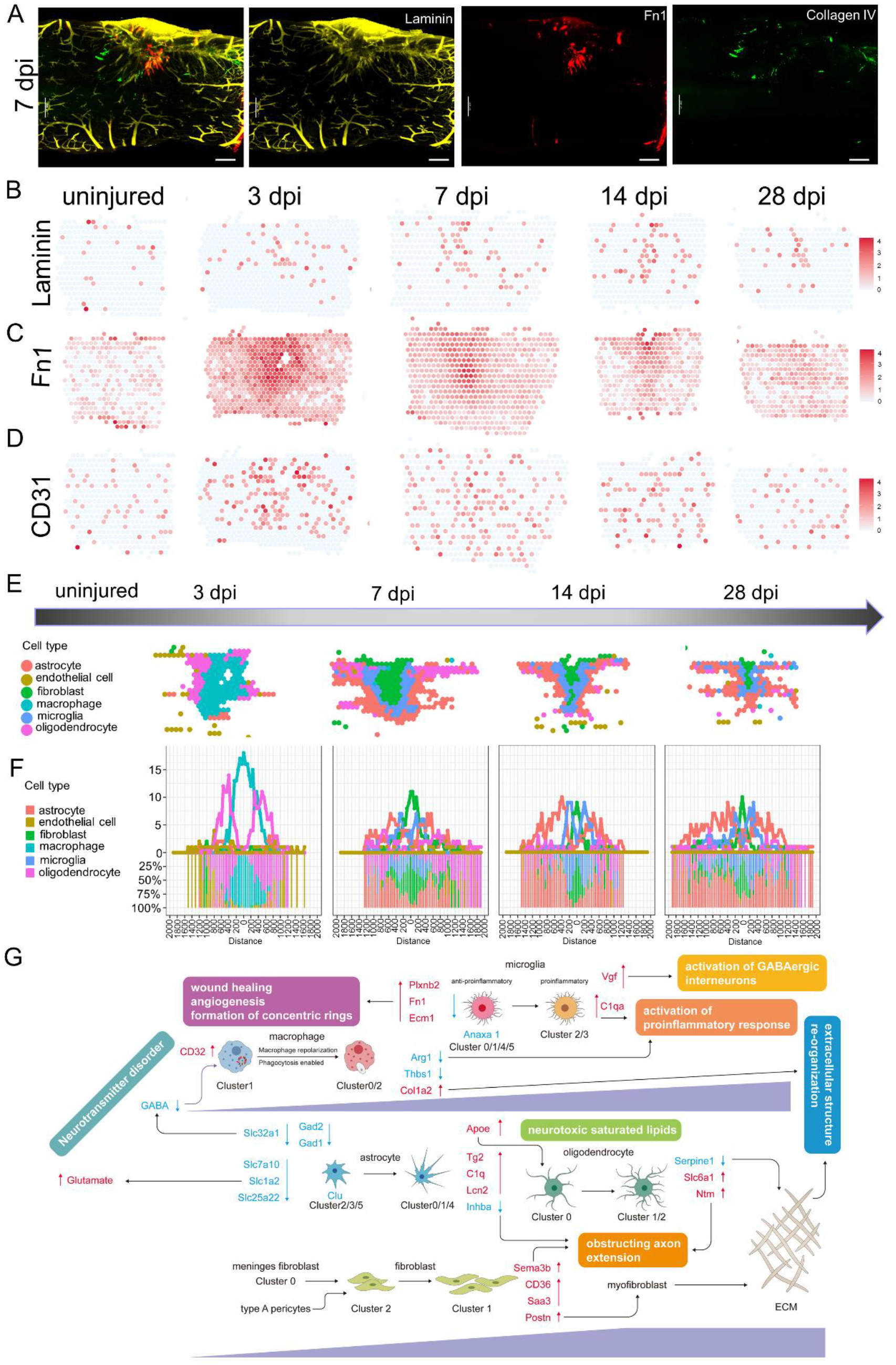
Visualization and re-interpretion of of the scar with transparency and ST. (A) IF stained 7 dpi scars for Laminin (yellow), Fn1 (red) and Collagen IV (green) showing the extent of scar boundary. Scare bar, 200 µm. (B) The spatial maps showing the expression of Laminin at different time points after injury. (C) The spatial maps showing the expression of Fn1 at different time points after injury. (D) The spatial maps showing the expression of CD31 at different time points after injury. (E) ST showing the extent of scar boundary of different cell type. (F) Quantification of the number and fraction of spots representing astrocyte, endothelial cell, fibroblast, macrophage, microglia and oligodendrocyte. (G) Summary of the changes in key genes and biological processes after SCI.

## Discussion

After the adult mammal’s CNS is injured, the scar tissue formed is essential to seal the damaged tissue and control the damage expansion, but it is also a barrier to nerve regeneration. A mature CNS scar is a compartmentalized structure that contains a variety of cells. For example, activated astrocytes surround the outer layer of fibrous scar tissue and constitute the glial component of the scar. Non-neural cells, including local and infiltrating immune cells and fibroblasts that produce extracellular matrix, gather in the core of the scar and constitute the fibrosis or matrix component of the scar. Although many different types of cells are thought to be involved in fibrotic scars, it is difficult to accurately track and observe their genetic fate in vivo due to the lack of a lineage tracing system for the corresponding cells. scRNA-seq allows us to use single-cell transcription profiles to study cellular heterogeneity in the CNS, thereby improving our understanding of CNS diseases. However, scRNA-seq generally do not involve spatial patterns of gene expression. Here, we provided a comprehensive resource map of the glial scars formed after spinal cord hemisection. For the first time, we constructed the measurement of gene expression in the mouse spinal cord tissue during the scar formation with ST technology, describing cell lineage trajectories and spatial heterogeneity that single-cell technology cannot achieve.

### New hints of glial scar components obtained by ST technique

A mature central nervous system scar is a compartmentalized structure that contains a variety of cells (*9*). Activated astrocytes and microglia surround the outer layer of fibrotic scar. Non-neural cells, including local and infiltrating immune cells and fibroblasts, gather in the core of the scar. Using ST technology, we found that activated oligodendrocytes surrounded the outer layer and involved in immune homeostasis. Even scar-resident oligodendrocytes obstructed axon extension through secreted Ntm. Additionally, the traditional view is that scar-resident macrophages come from the blood supply (*58*) (*64*). But recent studies have shown that monocytes after spinal cord injury can come from adjacent skull and spine bone marrow (*66*). Our ST diagrams also verified the majority of scar-resident macrophages were derived from adjacent skull in our model.

### Spatiotempora-specific action of glial scars in spinal cord repair deciphered by ST technique

The current general view of glial scars is that glial scars have detrimental properties (inhibiting axon regeneration) and beneficial properties (protecting spare nerve tissue), but their exact time-specific roles are still poorly understood (*18, 71*). Although scar-resident fibroblasts obstructed axon extension at intermediate stage (14 dpi, 28 dpi), they secreted Bmp7 to support the survival of scar-resident oligodendrocytes. During the process of gliosis and fibrosis, the bidirectional ligand-receptor interaction on the cluster boundary helps to maintain the scar structure. The spatiotemporal changes of the cell’s position in the scar may be related to its function. A special subset of neonatal microglia can form bridges between the two stumps after SCI and mediate the axon regeneration (*85*). Through ST analysis, we also found that a special subset of adult microglia can cross the fibrotic scar and build a bridge. But these bridges eventually disappeared at 28 dpi, suggesting the difference of regeneration ability at different stages of development is closely related to scar microenvironment. ST also identified that Cluster 1 cells were likely to be M2 macrophages and always in the core of injury site. Interesting, cluster 1 cells have high expression level of Ccr2, suggesting they were bone marrow-derived mononuclear macrophages.

### New treatment strategies to eliminate scars provided by ST technology

In recent years, the reconstruction of regenerative microenvironment based on biomaterials and the promotion of functional rehabilitation have become one of the frontiers of spinal cord injury treatment. The ideal scaffolds have good biocompatibility, biodegradability and biomechanical properties. These characteristics help to more effectively repair SCI through the reconstruction of regenerative microenvironment (*116*). For example, neurotrophin-3 (NT-3)-loaded chitosan biodegradable material (*117*) and small molecules combined with collagen hydrogel (*118*) enabled robust neural regeneration and functional restoration. To reduce microglia/macrophage mediated-inflammation, a combination of photo-crosslinked hydrogel transplantation and colony-stimulating factor 1 receptor (CSF1R) inhibitor (PLX3397) treatment inhibited pro-inflammatory factors via activated microglia/macrophages depletion and promoted endogenous neural stem/progenitor cells (NSPCs) neurogenesis (*119*). Spatial transcriptomics technologies provide an unbiased spatial composition map and gradually become a powerful tool to decipher tissue development and pathology (*120*). Except for microglia/macrophages, our ST found both astrocytes and oligodendrocytes involved in maintaining inflammatory homeostasis. Specially, the effects of oligodendrocytes are not well understood. CLU derived from plasma has been proved to be a pivotal anti-inflammatory effector in a mouse model of acute brain inflammation and a mouse model of Alzheimer’s disease (*105*). The CLU-rs9331896-TT genotype was even a risk factor for Parkinson’s disease (PD) (*121, 122*). ST showed activated astrocytes had high expression level of Clu at the acute stage and subacute stage, and gradually decreased at intermediate stage. CLU-loaded biodegradable material can control release rate and might help to reverse the deficiency of CLU at intermediate stage for glial scars treatment. Another issue is the persistent existence of CD36^+^ myofibroblast in the scar core. Tissue-resident fibroblasts have the known role in scar fibrosis (*123, 124*), and emerge as a key cell type in regulating activation or suppression of an immune response (*125*), For example, dermal fibroblasts that recruit and activate CD8^+^ cytotoxic T cells through secreted chemokines are responsible for driving patterned autoimmune activity in vitiligo (*126*). In our model, SCI induced a significant fibroblast response, which formed fibrotic scar that restrict the regeneration of adult mammalian axon. Both our ST and the previous study have proved that scar associated fibroblasts are derived from type A pericytes (*127*). We hypothesize that we can promote functional recovery by eliminating scar associated fibroblasts. Fortunately, a moderate reduction by genetic strategy in pericyte-derived fibrosis can reduce scar pathology and achieve functional recovery (*74*). ST analysis showed that CD36, which is involved in skin scar formation, was obviously activated in scar-resident fibroblasts. This suggests that CD36 may become a therapeutic target for spinal cord scars. Salvianolic acid, an inhibitor of CD36, may become a new therapy for drugs to eliminating scar-resident fibroblasts (*83, 128, 129*). Additionally, during skin wound healing in the mouse, adipocytes regenerate from myofibroblasts, a cell type thought to be differentiated and nonadipogenic (*130*), suggesting the possibility that scar-resident fibroblasts can be eliminating through the reprogramming (*131*).

Spatial transcriptomics maps of healthy or diseased tissues facilitate unbiased exploration and hypothesis generation. We found that the spatial mapping of the differentiation transition more comprehensively describes the possible trajectories between glial cells, fibroblasts and immune cell subpopulations, and more accurately defines the scar boundary and the extent of the lesion. Thus far, the current ST method still faces challenges, including resolution and sensitivity limitations, and throughput. But, with the continuous updating of technology, these problems are being gradually overcome by researchers.

Our transcriptome data set aims to present a comprehensive decoding of glial scars, and will serve as a reference for future research. We believe that our data will help understand the spatial gene expression and gene regulatory network of spinal glial scars, and provide ideas for potential clinical treatment strategies. Last, we present all aspects of these highly dimensional data via an interactive resource, spatio-temporal cell atlas of the mouse spinal cord scar (STASCS).

## Acknowledgement

We thank Lai Xu and Xiaoqing Yu for assistance in the preparation of the manuscript and figure. Thanks to the Shanghai Sinotech Genomics Co., Ltd. and Shanghai Biochip Co., Ltd. for excellent technical assistance.

---

**Figure. S1.**
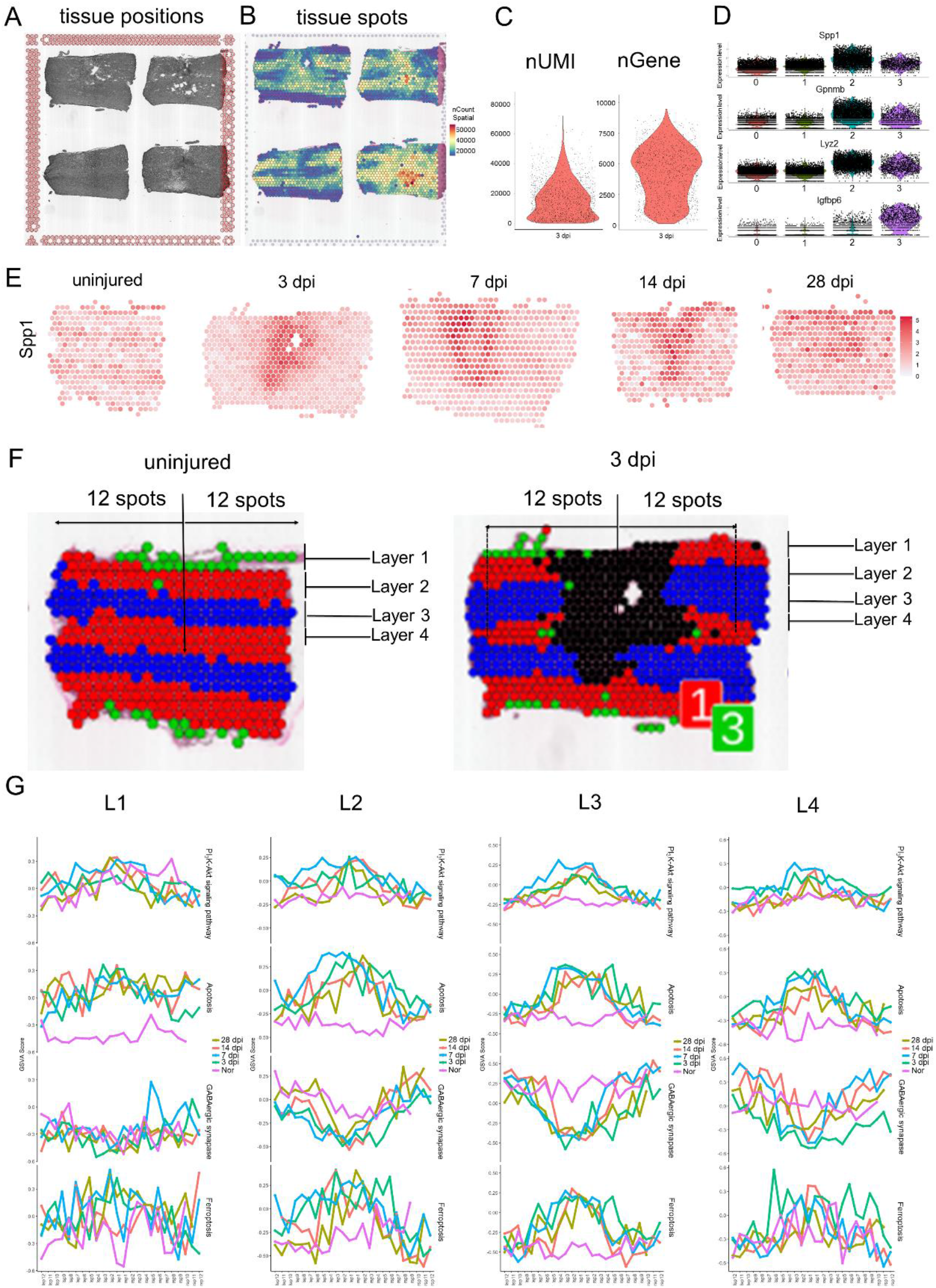
Quality control and spatial distribution of the spots involved in the glia scar formation. (A) The representative images of sections position from the samples at 3 dpi. (B) The representative images of the captured spots in the sections from the samples at 3 dpi. (C) Violin Plots showing the nUMI and nGene in the sections from the samples at 3 dpi. (D) Violin Plots showing the representative genes of cluster 2 cells and cluster 3 cells, the scar-associated clusters. (E) The spatial feature plots of Spp1, a stress response-associated-gene. (F) The symmetric distribution pattern of cluster 0 cells and cluster 1 cells in normal spinal cord, and these cells were divided into four layers from layer 1 to layer 4. Subsequently, these cells in the scar were divided into four layers by similar approaches. (G) Selected Gene Set Variation Analysis (GSVA) terms that were enriched in different layer at different stages of scar maturation. Fish’s exact test in the pathway enrichment was used.

**Figure. S2.**
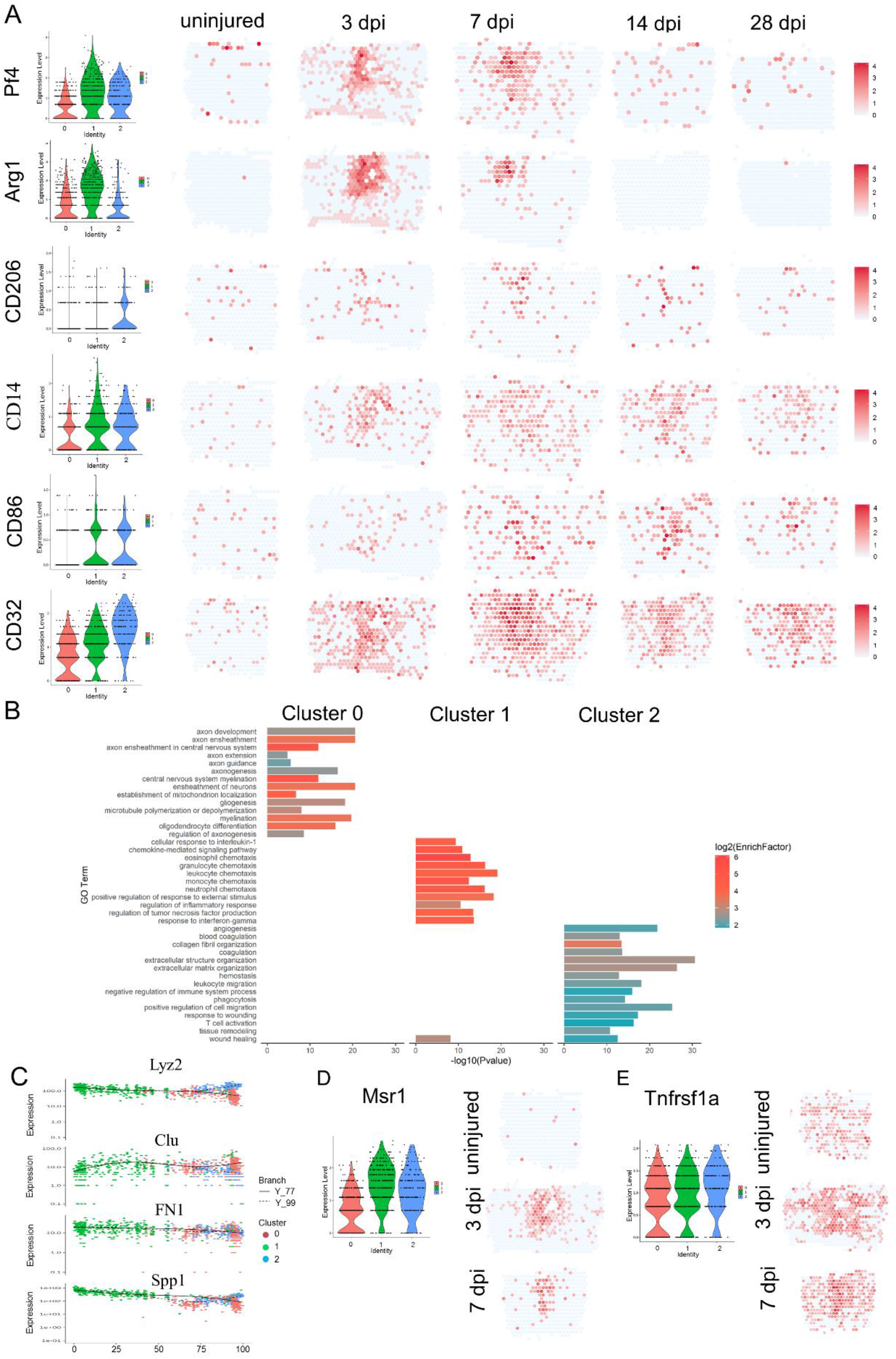
Distinct macrophage responses after lateral hemisection to spinal cord. (A) Violin Plots and the spatial maps showing the expression of marker genes at different time points after injury. (B) Selected GO terms that were enriched for each cluster. Fish’s exact test in the pathway enrichment was used. (C) The expression of selected genes correlating with pseudotime. (D and E) Violin Plots and the spatial maps showing the expression of selected genes at different time points after injury.

**Figure. S3.**
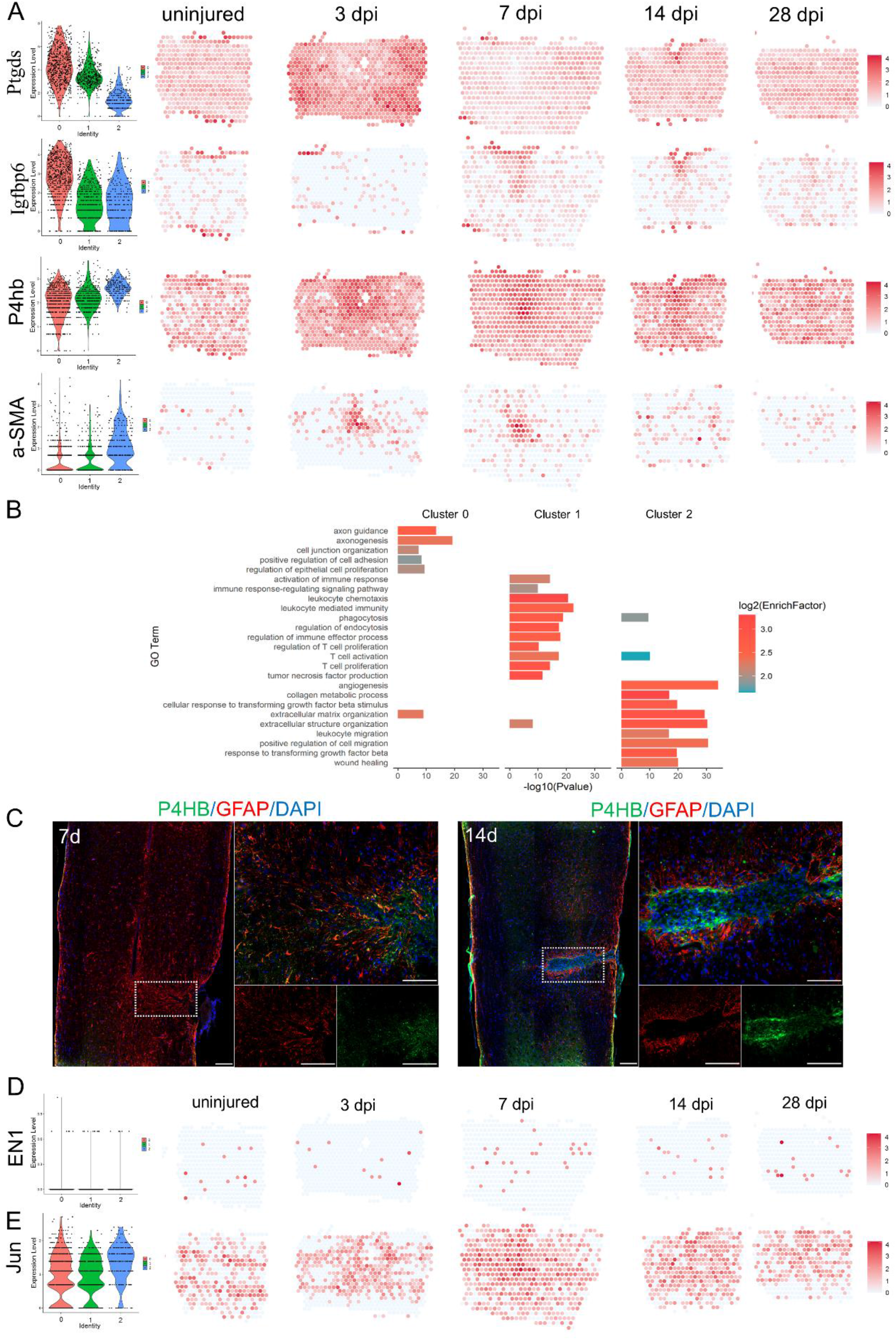
Distinct fibroblast responses after lateral hemisection to spinal cord. (A) Violin Plots and the spatial maps showing the expression of selected differentially expressed genes (DEGs) and marker genes at different time points after injury. (B) Selected GO terms that were enriched for each cluster. Fish’s exact test in the pathway enrichment was used. (C) IF stained 7 dpi and 14 dpi scars for GFAP (red), P4HB (green) and DAPI (blue) confirming the maturation of scar (n=3). Scare bar, 200 µm. (D) Violin Plots and the spatial maps showing the expression of En1 and Jun, two critical regulators of pathological skin scarring, at different time points after injury.

**Figure. S4.**
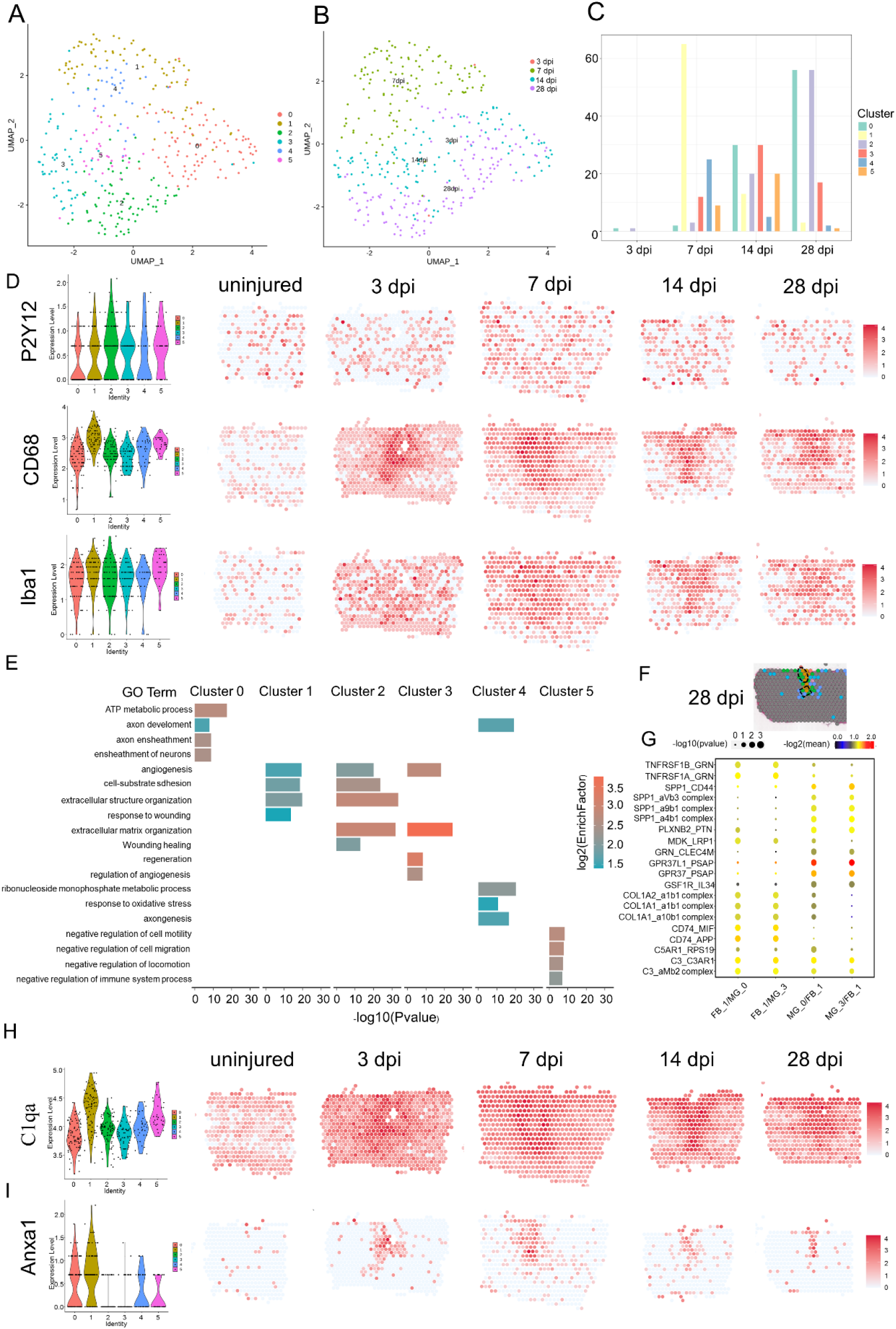
Distinct microglia responses after lateral hemisection to spinal cord. (A) UMAP spots showing the three clusters of microglia in the glia scar. (B) UMAP spots embedding overlay showing the distribution of spots at different time points after injury. (C) Histogram showing the number of spots in each subpopulation at different time points after injury. (D) Violin Plots and the spatial maps showing the expression of selected marker genes at different time points after injury. (E) Selected GO terms that were enriched for each cluster. Fish’s exact test in the pathway enrichment was used. (F) The definition of the boundary areas to study the interaction between two neighbor microglia clusters in the scar at 28 dpi. (G) Dot plot of the mean interaction scores between the neighbor clusters at 28 dpi. Size of the circle denoted p-value, and color denoted the mean interaction scores. (H) Violin Plots and the spatial maps showing the expression of C1q and Anxa1 at different time points after injury.

**Figure. S5.**
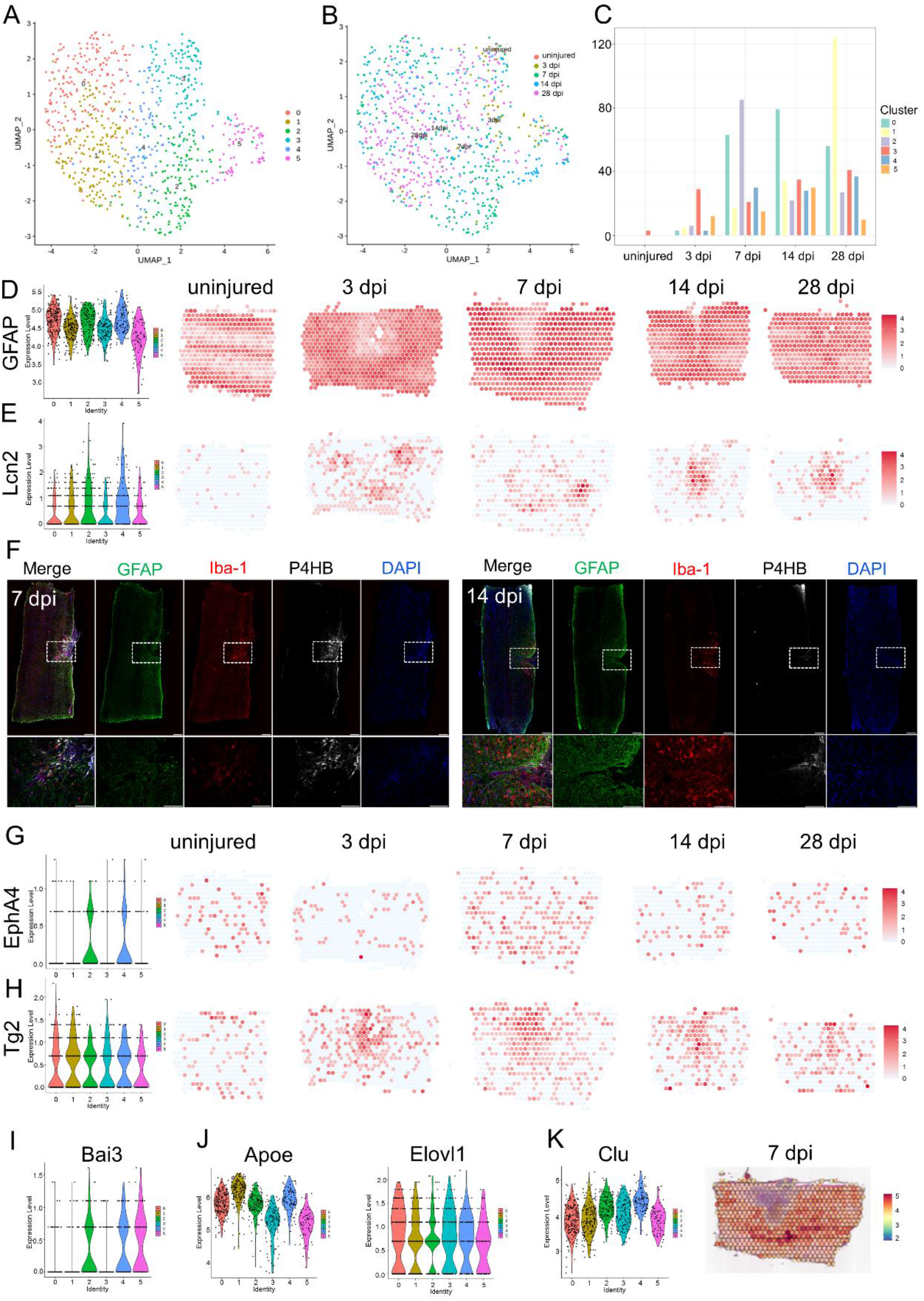
Distinct astrocyte responses after lateral hemisection to spinal cord. (A) UMAP spots showing the six clusters of astrocyte in the glia scar. (B) UMAP spots embedding overlay showing the distribution of spots at different time points after injury. (C) Histogram showing the number of spots in each subpopulation at different time points after injury. (D) Violin Plots and the spatial maps showing the expression of GFAP at different time points after injury. (E) Violin Plots and the spatial maps showing the expression of Lcn2 at different time points after injury. (F) IF stained 7 dpi and 14 dpi scars for GFAP (green), Iba1(red), P4HB (gray) and DAPI (blue) (n=3). Scare bar, 200 µm. (G) Violin Plots and the spatial maps showing the expression of EphA4 at different time points after injury. (H) Violin Plots and the spatial maps showing the expression of Tg2 at different time points after injury. (H) Violin Plots showing the expression of Bai3 in astrocytes scar after injury. (I) Violin Plots showing the expression of Apoe and Elovl1 in astrocytes scar after injury. (J) Violin Plots and the spatial map showing the expression of Clu in astrocytes scar after injury.

**Figure. S6.**
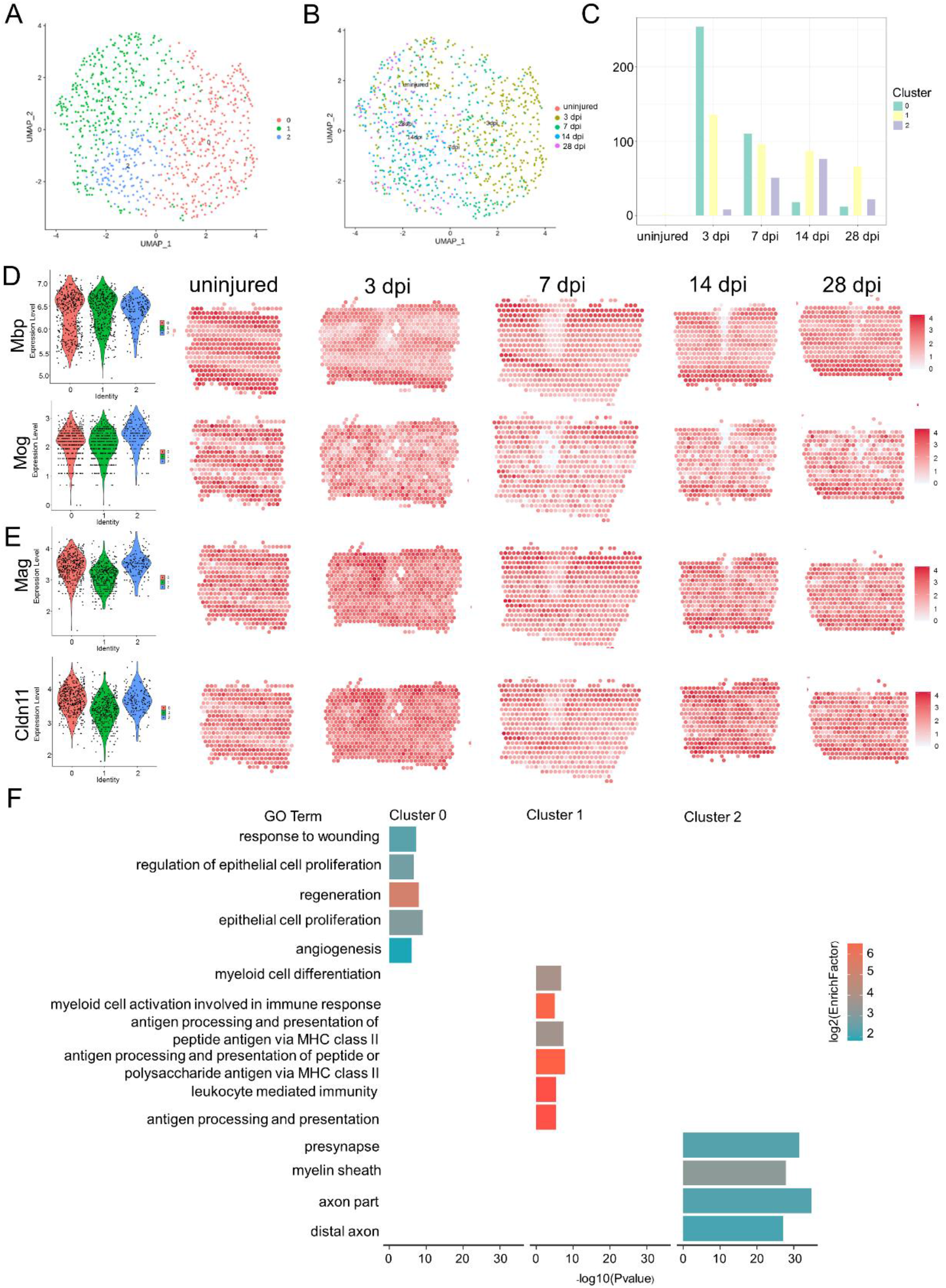
Distinct oligodendrocyte responses after lateral hemisection to spinal cord. (A) UMAP spots showing the three clusters of microglia in the glia scar. (B) UMAP spots embedding overlay showing the distribution of spots at different time points after injury. (C) Histogram showing the number of spots in each subpopulation at different time points after injury. (D) Violin Plots and the spatial maps showing the expression of Mbp and Mog at different time points after injury. (E) Violin Plots and the spatial maps showing the expression of Mag and Cldn11 at different time points after injury. (F) Selected GO terms that were enriched for each cluster. Fish’s exact test in the pathway enrichment was used.

**Figure. S7.**
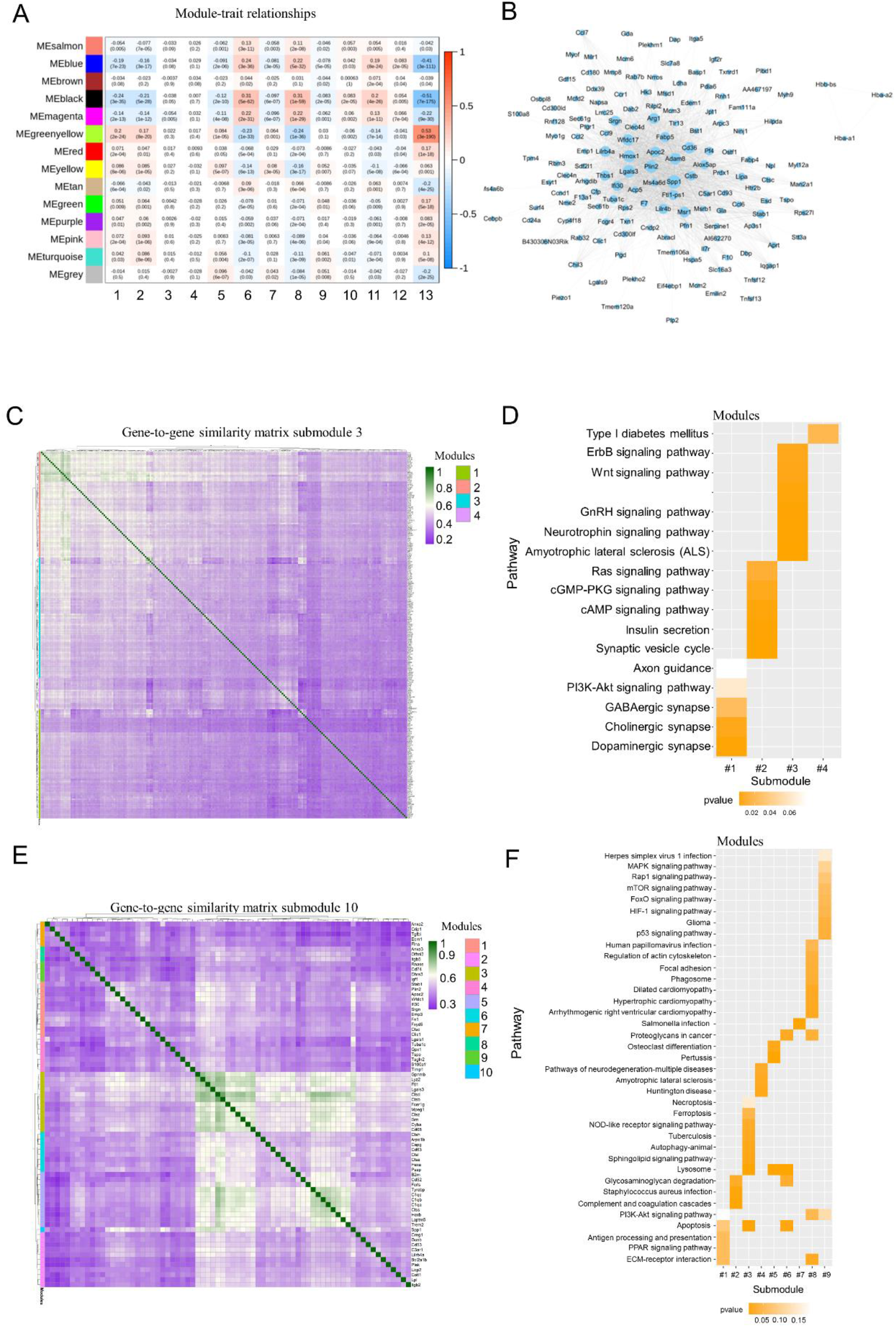
Spatiotemporal dynamics of genes expression during the maturation of scar. (A) The heatmap depicting the correlation scores (digit in the box above) as well as its corresponding P-value (digit in the box below) of modules (rows) and different celltype after injury (columns). 1, astrocyte; 2, microglia; 3, neuron and astrocyte; 4, ascular smooth muscle cell and endothelial cell; 5, fibroblast; 6, macrophage; 7, astrocyte; 8, oligodendrocyte; 9, astrocyte and oligodendrocyte; 10, endothelial cell; 11, macrophage; 12, fibroblast; 13, time. (B) Co-expression of selected hub genes enrichment in Meblack. The networks were created by Cytoscape. (C) Hierarchical clustering of gene in module 3. (D) Analysis of enriched KEGG pathway among the genes for submodules in (C). (E) Hierarchical clustering of gene in module 10. (F) Analysis of enriched KEGG pathway among the genes for submodules in (E).

## References

1. A. Alizadeh, S. M. Dyck, S. Karimi-Abdolrezaee, Traumatic Spinal Cord Injury: An Overview of Pathophysiology, Models and Acute Injury Mechanisms. Frontiers in neurology 10, 282 2019.

2. A. Anjum et al., Spinal Cord Injury: Pathophysiology, Multimolecular Interactions, and Underlying Recovery Mechanisms. International journal of molecular sciences 21, 2020.

3. H. Wang et al., Endocrine Therapy for the Functional Recovery of Spinal Cord Injury. Frontiers in neuroscience 14, 590570 2020.

4. L. L. Liau et al., Treatment of spinal cord injury with mesenchymal stem cells. Cell & Bioscience 10, 112 2020.

5. S. L. Lindsay, G. A. McCanney, A. G. Willison, S. C. Barnett, Multi-target approaches to CNS repair: olfactory mucosa-derived cells and heparan sulfates. Nature reviews. Neurology 16, 229–240 2020.

6. J. Silver, J. H. Miller, Regeneration beyond the glial scar. Nature reviews. Neuroscience 5, 146–156 2004.

7. Y. Ren et al., Ependymal cell contribution to scar formation after spinal cord injury is minimal, local and dependent on direct ependymal injury. Scientific reports 7, 41122 2017.

8. T. Yang, Y. Dai, G. Chen, S. Cui, Dissecting the Dual Role of the Glial Scar and Scar-Forming Astrocytes in Spinal Cord Injury. Frontiers in cellular neuroscience 14, 78 2020.

9. V. Bellver-Landete et al., Microglia are an essential component of the neuroprotective scar that forms after spinal cord injury. Nature communications 10, 518 2019.

10. I. B. Wanner et al., Glial scar borders are formed by newly proliferated, elongated astrocytes that interact to corral inflammatory and fibrotic cells via STAT3-dependent mechanisms after spinal cord injury. The Journal of neuroscience : the official journal of the Society for Neuroscience 33, 12870–12886 2013.

11. C. C. John Lin et al., Identification of diverse astrocyte populations and their malignant analogs. Nature neuroscience 20, 396–405 2017.

12. M. Y. Batiuk et al., Identification of region-specific astrocyte subtypes at single cell resolution. Nature communications 11, 1220 2020.

13. N. Habib et al., Disease-associated astrocytes in Alzheimer’s disease and aging. Nature neuroscience 23, 701–706 2020.

14. Y. Shi et al., Mouse and human share conserved transcriptional programs for interneuron development. Science (New York, N.Y.) 374, eabj6641 2021.

15. L. M. Milich et al., Single-cell analysis of the cellular heterogeneity and interactions in the injured mouse spinal cord. The Journal of experimental medicine 218, 2021.

16. R. Satija, J. A. Farrell, D. Gennert, A. F. Schier, A. Regev, Spatial reconstruction of single-cell gene expression data. Nature biotechnology 33, 495–502 2015.

17. K. Achim et al., High-throughput spatial mapping of single-cell RNA-seq data to tissue of origin. Nature biotechnology 33, 503–509 2015.

18. B. Yu et al., The Landscape of Gene Expression and Molecular Regulation Following Spinal Cord Hemisection in Rats. Frontiers in molecular neuroscience 12, 287 2019.

19. R. Ke et al., In situ sequencing for RNA analysis in preserved tissue and cells. Nature methods 10, 857–860 2013.

20. J. H. Lee et al., Highly multiplexed subcellular RNA sequencing in situ. Science (New York, N.Y.) 343, 1360–1363 2014.

21. C. L. Eng et al., Transcriptome-scale super-resolved imaging in tissues by RNA seqFISH. Nature 568, 235–239 2019.

22. X. Wang et al., Three-dimensional intact-tissue sequencing of single-cell transcriptional states. Science (New York, N.Y.) 361, 2018.

23. Y. Wang et al., EASI-FISH for thick tissue defines lateral hypothalamus spatio-molecular organization. Cell, 2021.

24. X. Qian et al., Probabilistic cell typing enables fine mapping of closely related cell types in situ. Nature methods 17, 101–106 2020.

25. P. L. Ståhl et al., Visualization and analysis of gene expression in tissue sections by spatial transcriptomics. Science (New York, N.Y.) 353, 78–82 2016.

26. S. Giacomello et al., Spatially resolved transcriptome profiling in model plant species. Nature plants 3, 17061 2017.

27. S. Giacomello, J. Lundeberg, Preparation of plant tissue to enable Spatial Transcriptomics profiling using barcoded microarrays. Nature Protocols 13, 2425–2446 2018.

28. A. Lundmark et al., Gene expression profiling of periodontitis-affected gingival tissue by spatial transcriptomics. Scientific reports 8, 9370 2018.

29. F. Salmén et al., Barcoded solid-phase RNA capture for Spatial Transcriptomics profiling in mammalian tissue sections. Nat Protoc 13, 2501–2534 2018.

30. C. Ortiz et al., Molecular atlas of the adult mouse brain. Science advances 6, eabb3446 2020.

31. S. Maniatis et al., Spatiotemporal dynamics of molecular pathology in amyotrophic lateral sclerosis. Science (New York, N.Y.) 364, 89–93 2019.

32. M. Asp et al., Spatial detection of fetal marker genes expressed at low level in adult human heart tissue. Scientific reports 7, 12941 2017.

33. M. Asp et al., A Spatiotemporal Organ-Wide Gene Expression and Cell Atlas of the Developing Human Heart. Cell 179, 1647–1660.e1619 2019.

34. R. van den Brand et al., Restoring Voluntary Control of Locomotion after Paralyzing Spinal Cord Injury. Science (New York, N.Y.) 336, 1182–1185 2012.

35. S. Hänzelmann, R. Castelo, J. Guinney, GSVA: gene set variation analysis for microarray and RNA-seq data. BMC bioinformatics 14, 7 2013.

36. X. J. Qiu et al., Reversed graph embedding resolves complex single-cell trajectories. Nature methods 14, 979-+ 2017.

37. P. Langfelder, S. Horvath, WGCNA: an R package for weighted correlation network analysis. BMC bioinformatics 9, 2008.

38. S. Maniatis et al., Spatiotemporal dynamics of molecular pathology in amyotrophic lateral sclerosis. Science (New York, N.Y.) 364, 89-+ 2019.

39. R. Vento-Tormo et al., Single-cell reconstruction of the early maternal-fetal interface in humans. Nature 563, 347-+ 2018.

40. T. Liebmann et al., Three-Dimensional Study of Alzheimer’s Disease Hallmarks Using the iDISCO Clearing Method. Cell reports 16, 1138–1152 2016.

41. B. Chen et al., Reactivation of Dormant Relay Pathways in Injured Spinal Cord by KCC2 Manipulations. Cell 174, 521-+ 2018.

42. M. A. Anderson et al., Astrocyte scar formation aids central nervous system axon regeneration. Nature 532, 195–200 2016.

43. C. Soderblom et al., Perivascular fibroblasts form the fibrotic scar after contusive spinal cord injury. The Journal of neuroscience : the official journal of the Society for Neuroscience 33, 13882–13887 2013.

44. K. X. Wang, D. T. Denhardt, Osteopontin: role in immune regulation and stress responses. Cytokine & growth factor reviews 19, 333–345 2008.

45. G. Cappellano et al., The Yin-Yang of osteopontin in nervous system diseases: damage versus repair. Neural regeneration research 16, 1131–1137 2021.

46. X. Duan et al., Subtype-specific regeneration of retinal ganglion cells following axotomy: effects of osteopontin and mTOR signaling. Neuron 85, 1244–1256 2015.

47. F. Kahles, H. M. Findeisen, D. Bruemmer, Osteopontin: A novel regulator at the cross roads of inflammation, obesity and diabetes. Molecular metabolism 3, 384–393 2014.

48. M. Hashimoto, D. Sun, S. R. Rittling, D. T. Denhardt, W. Young, Osteopontin-deficient mice exhibit less inflammation, greater tissue damage, and impaired locomotor recovery from spinal cord injury compared with wild-type controls. The Journal of neuroscience : the official journal of the Society for Neuroscience 27, 3603–3611 2007.

49. M. Ahmed, G. C. Kundu, Osteopontin selectively regulates p70S6K/mTOR phosphorylation leading to NF-kappaB dependent AP-1-mediated ICAM-1 expression in breast cancer cells. Molecular cancer 9, 101 2010.

50. M. C. Wright et al., Novel roles for osteopontin and clusterin in peripheral motor and sensory axon regeneration. The Journal of neuroscience : the official journal of the Society for Neuroscience 34, 1689–1700 2014.

51. R. Rahimian, L. C. Beland, J. Kriz, Galectin-3: mediator of microglia responses in injured brain. Drug Discov Today 23, 375–381 2018.

52. M. Puigdellivol, D. H. Allendorf, G. C. Brown, Sialylation and Galectin-3 in Microglia-Mediated Neuroinflammation and Neurodegeneration. Frontiers in cellular neuroscience 14, 2020.

53. M. A. DePaul, C. Y. Lin, J. Silver, Y. S. Lee, Peripheral Nerve Transplantation Combined with Acidic Fibroblast Growth Factor and Chondroitinase Induces Regeneration and Improves Urinary Function in Complete Spinal Cord Transected Adult Mice. PloS one 10, e0139335 2015.

54. C. Göritz et al., A pericyte origin of spinal cord scar tissue. Science (New York, N.Y.) 333, 238–242 2011.

55. Y. Zhang et al., Ferroptosis inhibitor SRS 16-86 attenuates ferroptosis and promotes functional recovery in contusion spinal cord injury. Brain Res 1706, 48–57 2019.

56. G. Courtine, M. V. Sofroniew, Spinal cord repair: advances in biology and technology. Nature medicine 25, 898–908 2019.

57. W. Xie et al., Pterostilbene accelerates wound healing by modulating diabetes-induced estrogen receptor β suppression in hematopoietic stem cells. Burns & trauma 9, tkaa045 2021.

58. K. A. Kigerl et al., Identification of two distinct macrophage subsets with divergent effects causing either neurotoxicity or regeneration in the injured mouse spinal cord. The Journal of neuroscience : the official journal of the Society for Neuroscience 29, 13435–13444 2009.

59. R. L. Benton et al., Transcriptomic screening of microvascular endothelial cells implicates novel molecular regulators of vascular dysfunction after spinal cord injury. Journal of cerebral blood flow and metabolism : official journal of the International Society of Cerebral Blood Flow and Metabolism 28, 1771–1785 2008.

60. Y. Pu, K. Meng, C. Gu, L. Wang, X. Zhang, Thrombospondin-1 modified bone marrow mesenchymal stem cells (BMSCs) promote neurite outgrowth and functional recovery in rats with spinal cord injury. Oncotarget 8, 96276–96289 2017.

61. M. Hara et al., Interaction of reactive astrocytes with type I collagen induces astrocytic scar formation through the integrin-N-cadherin pathway after spinal cord injury. Nature medicine 23, 818–828 2017.

62. F. Q. Kong et al., Macrophage MSR1 promotes the formation of foamy macrophage and neuronal apoptosis after spinal cord injury. J Neuroinflamm 17, 2020.

63. L. Cavone et al., A unique macrophage subpopulation signals directly to progenitor cells to promote regenerative neurogenesis in the zebrafish spinal cord. Dev Cell 56, 1617-+ 2021.

64. N. Cox et al., Diet-regulated production of PDGFcc by macrophages controls energy storage. Science (New York, N.Y.) 373, 2021.

65. E. I. Paschalis et al., Permanent neuroglial remodeling of the retina following infiltration of CSF1R inhibition-resistant peripheral monocytes. P Natl Acad Sci USA 115, E11359–E11368 2018.

66. A. Cugurra et al., Skull and vertebral bone marrow are myeloid cell reservoirs for the meninges and CNS parenchyma. Science (New York, N.Y.) 373, 409-+ 2021.

67. Y. Zhang et al., Deciphering glial scar after spinal cord injury. Burns & trauma 9, tkab035 2021.

68. E. Bellei et al., Serum protein changes in a rat model of chronic pain show a correlation between animal and humans. Scientific reports 7, 2017.

69. L. Micutkova et al., Insulin-like growth factor binding protein-6 delays replicative senescence of human fibroblasts. Mech Ageing Dev 132, 468–479 2011.

70. S. Wang et al., The Expression of IGFBP6 after Spinal Cord Injury: Implications for Neuronal Apoptosis. Neurochem Res 42, 455–467 2017.

71. K. L. Adams, V. Gallo, The diversity and disparity of the glial scar. Nature neuroscience 21, 9–15 2018.

72. D. O. Dias et al., Pericyte-derived fibrotic scarring is conserved across diverse central nervous system lesions. Nature communications 12, 2021.

73. C. Goritz et al., A Pericyte Origin of Spinal Cord Scar Tissue. Science (New York, N.Y.) 333, 238–242 2011.

74. D. O. Dias et al., Reducing Pericyte-Derived Scarring Promotes Recovery after Spinal Cord Injury. Cell 173, 153-+ 2018.

75. C. Kuppe et al., Decoding myofibroblast origins in human kidney fibrosis. Nature 589, 281-+ 2021.

76. H. Wu et al., Periostin expression induced by oxidative stress contributes to myocardial fibrosis in a rat model of high salt-induced hypertension. Mol Med Rep 14, 776–782 2016.

77. K. Izuhara et al., Periostin in inflammation and allergy. Cell Mol Life Sci 74, 4293–4303 2017.

78. C. H. Shih, M. Lacagnina, K. Leuer-Bisciotti, C. Proschel, Astroglial-Derived Periostin Promotes Axonal Regeneration after Spinal Cord Injury. Journal of Neuroscience 34, 2438–2443 2014.

79. K. Yokota et al., Periostin Promotes Scar Formation through the Interaction between Pericytes and Infiltrating Monocytes/Macrophages after Spinal Cord Injury. Am J Pathol 187, 639–653 2017.

80. P. Puthanveetil, S. L. Chen, B. A. Feng, A. Gautam, S. Chakrabarti, Long non-coding RNA MALAT1 regulates hyperglycaemia induced inflammatory process in the endothelial cells. J Cell Mol Med 19, 1418–1425 2015.

81. M. Djurec et al., Saa3 is a key mediator of the protumorigenic properties of cancer-associated fibroblasts in pancreatic tumors. P Natl Acad Sci USA 115, E1147–E1156 2018.

82. S. Mascharak et al., Preventing Engrailed-1 activation in fibroblasts yields wound regeneration without scarring. Science (New York, N.Y.) 372, 362-+ 2021.

83. M. F. Griffin et al., JUN promotes hypertrophic skin scarring via CD36 in preclinical in vitro and in vivo models. Sci Transl Med 13, 2021.

84. S. A. Myers, K. R. Andres, T. Hagg, S. R. Whittemore, CD36 deletion improves recovery from spinal cord injury. Exp Neurol 256, 25–38 2014.

85. Y. Li et al., Microglia-organized scar-free spinal cord repair in neonatal mice. Nature 587, 613-+ 2020.

86. E. Van Battum et al., Plexin-B2 controls the timing of differentiation and the motility of cerebellar granule neurons. Elife 10, 2021.

87. X. Zhou et al., Microglia and macrophages promote corralling, wound compaction and recovery after spinal cord injury via Plexin-B2. Nature neuroscience 23, 337-+ 2020.

88. R. C. Meyer, M. M. Giddens, B. M. Coleman, R. A. Hall, The protective role of prosaposin and its receptors in the nervous system. Brain Res 1585, 1–12 2014.

89. M. M. T. van Leent et al., Prosaposin mediates inflammation in atherosclerosis. Sci Transl Med 13, 2021.

90. P. Ciana et al., The orphan receptor GPR17 identified as a new dual uracil nucleotides/cysteinyl-leukotrienes receptor. The EMBO journal 25, 4615–4627 2006.

91. S. Ceruti et al., The P2Y-like receptor GPR17 as a sensor of damage and a new potential target in spinal cord injury. Brain 132, 2206–2218 2009.

92. D. Lecca et al., The Recently Identified P2Y-Like Receptor GPR17 Is a Sensor of Brain Damage and a New Target for Brain Repair. PloS one 3, 2008.

93. J. Wang et al., Robust Myelination of Regenerated Axons Induced by Combined Manipulations of GPR17 and Microglia. Neuron 108, 876-+ 2020.

94. S. A. Liddelow et al., Neurotoxic reactive astrocytes are induced by activated microglia. Nature 541, 481–487 2017.

95. Y. Goldshmit, S. McLenachan, A. Turnley, Roles of Eph receptors and ephrins in the normal and damaged adult CNS. Brain Res Rev 52, 327–345 2006.

96. X. Z. Yu, J. Nagai, B. S. Khakh, Improved tools to study astrocytes. Nature Reviews Neuroscience 21, 121–138 2020.

97. P. Bazzigaluppi, I. Weisspapir, B. Stefanovic, L. Leybaert, P. L. Carlen, Astrocytic gap junction blockade markedly increases extracellular potassium without causing seizures in the mouse neocortex. Neurobiol Dis 101, 1–7 2017.

98. K. I. Tanaka et al., Involvement of SAPK/JNK Signaling Pathway in Copper Enhanced Zinc-Induced Neuronal Cell Death. Toxicol Sci 169, 293–302 2019.

99. T. Omura et al., Robust Axonal Regeneration Occurs in the Injured CAST/Ei Mouse CNS. Neuron 86, 1215–1227 2015.

100. S. Aibar et al., SCENIC: single-cell regulatory network inference and clustering. Nature methods 14, 1083-+ 2017.

101. J. S. Yang et al., Ephrin-A3 reverse signaling regulates hippocampal neuronal damage and astrocytic glutamate transport after transient global ischemia. J Neurochem 131, 383–394 2014.

102. A. Elahi et al., Deletion or Inhibition of Astrocytic Transglutaminase 2 Promotes Functional Recovery after Spinal Cord Injury. Cells 10, 2021.

103. J. Wang et al., RTN4/NoGo-receptor binding to BAI adhesion-GPCRs regulates neuronal development. Cell 184, 5869–5885.e5825 2021.

104. K. A. Guttenplan et al., Neurotoxic reactive astrocytes induce cell death via saturated lipids. Nature 599, 102-+ 2021.

105. Z. De Miguel et al., Exercise plasma boosts memory and dampens brain inflammation via clusterin. Nature, 2021.

106. A. M. Falcao et al., Disease-specific oligodendrocyte lineage cells arise in multiple sclerosis. Nature medicine 24, 1837-+ 2018.

107. K. Singh et al., The combined impact of IgLON family proteins Lsamp and Neurotrimin on developing neurons and behavioral profiles in mouse. Brain Res Bull 140, 5–18 2018.

108. F. B. Berry et al., FOXC1 is required for cell viability and resistance to oxidative stress in the eye through the transcriptional regulation of FOXO1A. Hum Mol Genet 17, 490–505 2008.

109. D. Y. He et al., Chd7 cooperates with Sox10 and regulates the onset of CNS myelination and remyelination. Nature neuroscience 19, 678-+ 2016.

110. X. Wang, J. M. Xu, Y. P. Wang, L. Yang, Z. J. Li, Protective effects of BMP-7 against tumor necrosis factor α-induced oligodendrocyte apoptosis. International journal of developmental neuroscience : the official journal of the International Society for Developmental Neuroscience 53, 10–17 2016.

111. S. X. Liu et al., Overexpression of bone morphogenetic protein 7 reduces oligodendrocytes loss and promotes functional recovery after spinal cord injury. J Cell Mol Med 25, 8764–8774 2021.

112. H. Su, N. Na, X. Zhang, Y. Zhao, The biological function and significance of CD74 in immune diseases. Inflamm Res 66, 209–216 2017.

113. C. Zhao et al., Diffusion tensor imaging of spinal cord parenchyma lesion in rat with chronic spinal cord injury. Magn Reson Imaging 47, 25–32 2018.

114. J. S. Rao et al., Image correction for diffusion tensor imaging of Rhesus monkey thoracic spinal cord. J Med Primatol 48, 320–328 2019.

115. V. Bellver-Landete et al., Microglia are an essential component of the neuroprotective scar that forms after spinal cord injury. Nature communications 10, 2019.

116. X. Gu, Biodegradable Materials and the Tissue Engineering of Nerves. Engineering, 2021.

117. J. S. Rao et al., NT3-chitosan enables de novo regeneration and functional recovery in monkeys after spinal cord injury. P Natl Acad Sci USA 115, E5595–E5604 2018.

118. Y. M. Yang et al., Small molecules combined with collagen hydrogel direct neurogenesis and migration of neural stem cells after spinal cord injury. Biomaterials 269, 2021.

119. D. Z. Ma et al., A novel hydrogel-based treatment for complete transection spinal cord injury repair is driven by microglia/macrophages repopulation. Biomaterials 237, 2020.

120. A. Rao, D. Barkley, G. S. Franca, I. Yanai, Exploring tissue architecture using spatial transcriptomics. Nature 596, 211–220 2021.

121. Y. W. Lin et al., Association of CLU gene polymorphism with Parkinson’s disease in the Chinese Han population. J Gene Med 23, 2021.

122. M. S. Uddin et al., Exploring the Role of CLU in the Pathogenesis of Alzheimer’s Disease (Aug, 10.1007/s12640-020-00271-4, 2020). Neurotox Res 38, 1062–1062 2020.

123. N. C. Henderson, F. Rieder, T. A. Wynn, Fibrosis: from mechanisms to medicines. Nature 587, 555–566 2020.

124. M. V. Plikus et al., Fibroblasts: Origins, definitions, and functions in health and disease. Cell 184, 3852–3872 2021.

125. S. Davidson et al., Fibroblasts as immune regulators in infection, inflammation and cancer. Nat Rev Immunol 21, 704–717 2021.

126. Z. Xu et al., Anatomically distinct fibroblast subsets determine skin autoimmune patterns. Nature, 2021.

127. E. Llorens-Bobadilla et al., A latent lineage potential in resident neural stem cells enables spinal cord repair. Science (New York, N.Y.) 370, 73-+ 2020.

128. J. Yang, K. W. Park, S. Cho, Inhibition of the CD36 receptor reduces visceral fat accumulation and improves insulin resistance in obese mice carrying the BDNF-Val66Met variant. The Journal of biological chemistry 293, 13338–13348 2018.

129. Y. Bao et al., Salvianolic acid B inhibits macrophage uptake of modified low density lipoprotein (mLDL) in a scavenger receptor CD36-dependent manner. Atherosclerosis 223, 152–159 2012.

130. M. V. Plikus et al., Regeneration of fat cells from myofibroblasts during wound healing. Science (New York, N.Y.) 355, 748-+ 2017.

131. H. Qin, A. D. Zhao, K. Ma, X. B. Fu, Chemical conversion of human and mouse fibroblasts into motor neurons. Sci China Life Sci 61, 1151–1167 2018.

